# S1PR3 mediates glial stimulated tumor invasion in response to interstitial fluid flow

**DOI:** 10.64898/2025.12.01.690522

**Authors:** R. Chase Cornelison, Samantha Howerton, Kinsley M Tate, Caleb A. Stine, Alexis Petrosky, Kathryn M. Kingsmore, Sharon K. Michelhaugh, Jessica J. Cunningham, Katie Degen, Austin Hoggarth, Cora Carman-Esparza, Peng Jin, Maosen Wang, Caroline O’Brien, Aizhen Xiao, Benjamin W. Purow, James W. Mandell, Ian F. Kimbrough, Mark R. Witcher, Jennifer M. Munson

**Affiliations:** Fralin Biomedical Research Institute at Virginia Tech-Carilion; Department of Biomedical Engineering, Virginia Tech; Department of Neurosurgery, Carilion School of Medicine; Department of Biomedical Engineering, University of Massachussetts-Amherst; Department of Neuroscience, University of Virginia; Department of Pathology, University of Virginia; Department of Neurology, University of Virginia; Department of Biomedical Engineering, University of Virginia

## Abstract

Cellular invasion is a primary challenge to complete resection and treatment of glioblastoma, the most aggressive and deadly primary brain tumor. The brain tumor microenvironment actively stimulates glioma invasion through a multitude of cellular, chemical, and biophysical cues. We and others have shown elevated interstitial fluid flow at the tumor border is one such biophysical cue that directly stimulates invasion through tumor-intrinsic signaling and, in other tumor types, priming of cancer-associated stromal cells. It is currently unclear if interstitial flow similarly primes neuroglial cells to promote glioma cell dissemination and can be targeted for therapeutic purposes. Here, we show elevated interstitial flow upregulates expression of sphingosine-1-phosphate receptor 3 (S1PR3) in glial astrocytes and microglia, which drives glioma cell invasion via chemotaxis. Flow-induced expression of glial S1PR3 is tumor-independent and displays a biphasic relationship to fluid shear stress magnitude *in vitro* and flow rate *in vivo*. Inhibition of glial S1PR3 in a tissue engineered culture model and orthotopic mouse model abrogates flow-stimulated invasion, demonstrating a tumor-extrinsic approach to limiting glioblastoma progression. Given prior evidence of a pro-inflammatory role for glial S1PR3, identification of S1PR3 as a disease-agnostic marker of flow-stimulated glia may also have therapeutic implications across myriad neuropathologies.

## Introduction

Glioblastoma (GBM) is the most common malignant tumor in the central nervous system^1^ with a 5-year overall survival rate of 13% with standard of care^2^. Glioma cells are intrinsically plastic, acquiring diverse phenotypic states that support survival and invasion^3–5^. Phenotypic plasticity drives variable sensitivity to tumor-targeted therapies and selection toward drug-resistant states^4^. This issue is compounded by recent work identifying that these plastic cells are not a restricted subset of tumor cells but instead a state that most cells can adopt^6^. Thus, a major emphasis of emergent research for therapies against glioblastoma is how cues in the tumor microenvironment (TME) influence progression as targets less prone to plasticity.

Invasion into the healthy parenchyma drives tumor recurrence and follows distinct patterns related to bulk interstitial fluid flow within the brain^7^. This interstitial flow is increased in tumor-adjacent regions^8^ due to an elevated pressure differential between the tumor and surrounding parenchyma^9^. Pathological flow yields molecular responses that influence tumor cell invasion across cancers, with broad categories including mechanotransduction^10,11^ and autologous chemotaxis^12,13^. Most studies have focused on tumor-cell intrinsic processes stimulated by flow, but interstitial flow also primes tumor-associated stroma in cancers^9^. Previously, we identified roles for CXCR4/CXCL12-dependent autologous chemotaxis and CD44 in flow-enhanced glioma cell invasion^8,14^. However, the response is heterogeneous across patients, necessitating a search for additional targets^14^. Among these possible targets are responses driven by astrocytes and microglia, the most common glial subpopulations in the brain consistently implicated in GBM invasion and poor prognosis^15,16^. We sought to assess whether TME (i.e., glial-driven) mechanisms similarly mediate flow-enhanced glioma invasion and can be used to identify targets that are amenable to therapeutic development.

Here, we identify a role for glial sphingosine-1-phosphate receptor 3 (S1PR3) regulating flow-enhanced glioma invasion. We demonstrate peritumoral elevation of glial S1PR3 in high-flow regions across model systems that we verify experimentally and correlate with invasion. Leveraging our tissue engineered model of the patient TME^5,17^, we demonstrate efficacy of S1PR3 targeting across multiple patient-derived glioma cell lines that is dependent on the TME, and confirm efficacy in *vivo*.

## Results

### S1PR3 expression is increased in peritumoral regions of outward interstitial flow

Towards identifying molecular targets upregulated in flow regions within the TME, we leveraged our method to map interstitial fluid flow in live animals using dynamic contrast enhanced-magnetic resonance imaging (MRI) coupled with computational transport analysis^18,19^. The flow maps generated by MRI are validated to spatially coincide with extravasation of Evans Blue dye after intravenous delivery, which we used as a visual marker to identify and isolate tissue samples *ex vivo* for further analysis. We generated three patient-derived xenograft models using glioma stem cell lines G2, G34, and G528^20^ and confirmed tumor establishment by T2-weighted MRI. We then used computational analysis of dynamic contrast enhanced MRI analysis and Evans Blue fluorescence signal to identify peritumoral regions of high and low flow for gross dissection and analysis (Fig. 1a-c; Extended Fig. 1a). We also collected samples of the tumor bulk and “normal” tumor-free hemisphere as controls. Using microarray analysis, we identified nine genes across the three patient-derived xenograft models with the highest ratio of expression in “high” flow regions compared to low flow and normal tissue (Fig. 1d; Extended Fig. 1b-c). Of these genes, we selected sphingosine-1-phosphate receptor 3 (S1PR3) for further study, given the established role of this and other S1PRs in fluid mechanosensing^21^. We validated S1PR3 upregulation in regions of high flow relative to low flow by immunohistochemistry in the xenograft models (Fig. 1e).

**Figure 1.**
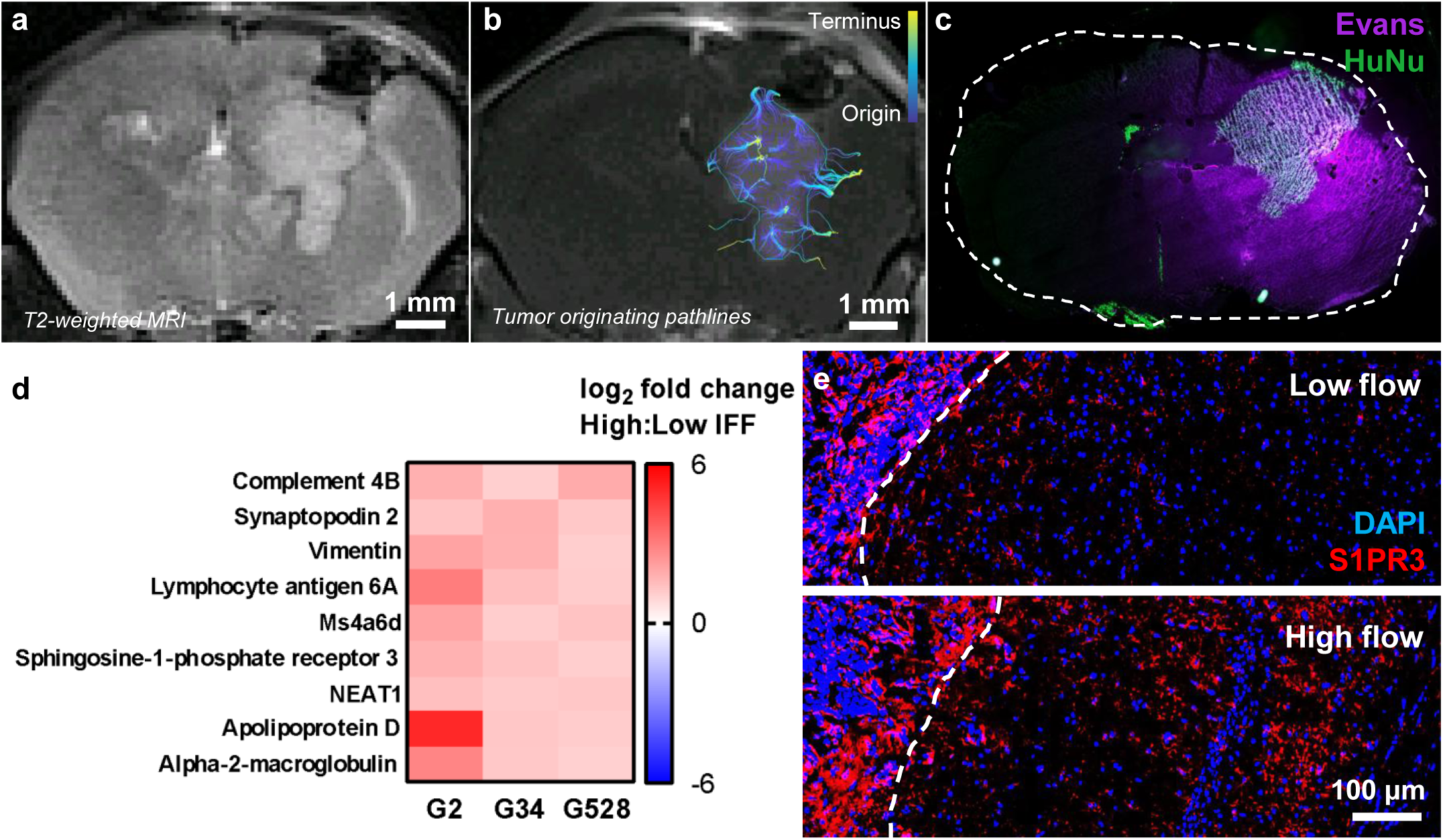
S1PR3 expression is elevated in regions of peritumoral flow relative to low-flow regions. **a.** Patient derived xenograft G2 imaged on T2 MRI 11 days post implantation. **b.** Tumor-originating pathlines as computationally calculated from Dynamic contrast enhanced MRI imaging on same day. **c.** Evans blue (magenta) delivered intravascularly on same day shown on histological section of tumor with tumor cells labeled by human nuclear antigen (HuNu) staining (green). **d.** Tissue was extracted from Evans blue-indicated high and low flow regions adjacent to the tumor and then analyzed by ArrayST 2.0 microarray. Top upregulated genes are shown across three different PDX models (n=3 mice per group for analysis) **e.** Immunohistochemistry for S1PR3 expression in identified high vs. low flow regions in G34 implanted PDX.

We then leveraged a mouse GL261 tumor model in the S1P_3_ ^mCherry^ reporter mouse line (B6.Cg-S1pr3tm1.1Hrose/J), which expresses the mCherry protein under the S1PR3 promoter^22^, to better visualize and spatially analyze S1PR3 upregulation specifically in peritumoral regions of flow. In combination, we also used an advanced MRI analysis algorithm capable of mapping flow pathlines originating from the tumor (Fig. 2a-e), which we have shown to capture nearly all invaded tumor cells within implanted tumor models and predict disease progression in mice^19^. We find that, in the majority of mice, mCherry fluorescence intensity (Fig. 2f; Extended Fig. 2a) significantly positively correlates with fluid velocity magnitude and density of tumor originating pathlines, whereas it negatively correlates with the distance to tumor boundary (Fig. 2g). Thus, reporter expression is highest near the tumor border, particularly regions where flow is leaving the tumor at a higher speeds.

### Glial cell S1PR3 correlates with peritumoral flow

We sought to identify which peritumoral cells express S1PR3. Using immunohistochemistry in the GL261-tumor bearing S1P3^mCherry^ reporter model, we find that mCherry expression primarily labels GLT-1+ or GFAP+ cells, representing astrocytes (Fig. 2h; Extended Fig. 2b). mCherry did not appear to label microglia (Fig. 2i), vascular endothelial cells (Extended Fig. 2c), pericytes (Extended Fig. 2d), or oligodendrocytes/precursor cells (Extended Fig. 2e-f).

**Figure 2.**
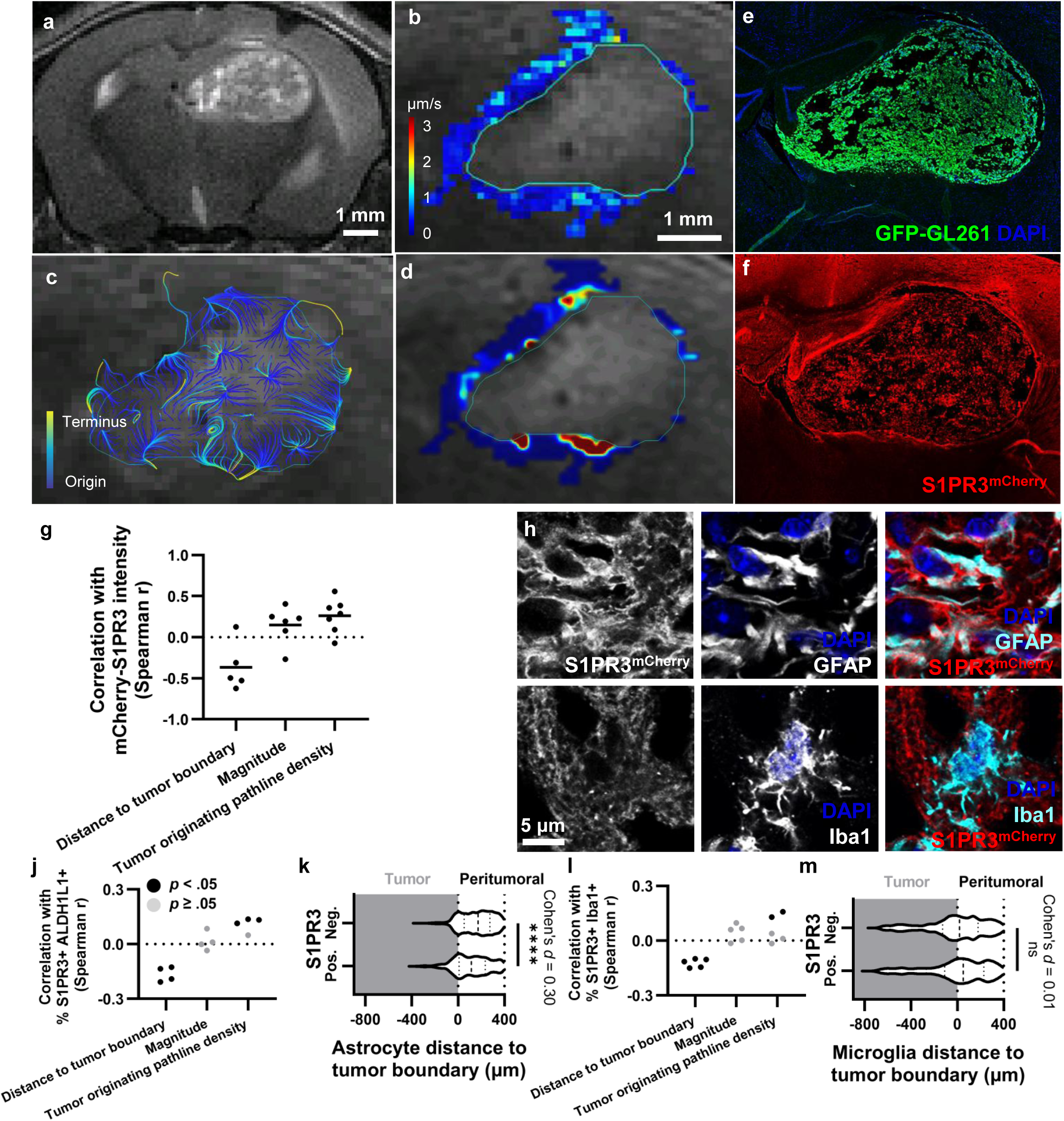
Peritumoral S1PR3 is enhanced in regions of outward flow in astrocytes. **a.** T2-weighted MRI of implanted GFP-GL261 tumors in S1PR3^mCherry^ mice at 15 days post implant. **b.** Interstitial velocity magnitude generated from DCE-MRI using Lymph4D analysis. **c.** tumor originating pathlines based on interstitial velocity vector fields indicating trajectories of flow from the tumor into the parenchyma. **d.** density of tumor originating pathlines that terminate in the parenchyma. **e.** GFP-GL261 (green) with DAPI (blue) on histological sections taken one day post DCE-MRI. **f.** S1PR3 reporter fluorescence on histological sections in the same mouse. **g.** Spatial correlation (Spearman r) of S1PR3^mCherry^ intensity with distance to tumor boundary, velocity magnitude, and tumor originating pathline density for each mouse. All datapoints shown are statistically significant (p < 0.05), n = 7 total mice analyzed. **h.** Representative immunohistochemical staining for S1PR3^mCherry^ (red) activated astrocytes (Cyan, GFAP), and DAPI (blue) in the parenchyma. **i.** Representative immunohistochemical staining for S1PR3^mCherry^ (red) microglia (Cyan, Iba1), and DAPI (blue) in the parenchyma. **j.** Classified S1PR3+ ALDH1L1+ cells (astrocytes) in parenchyma correlated with interstitial transport metrics (n = 4 mice). **k.** Classified ALDH1L1+ cells (astrocytes) by S1PR3 expression with distribution relative to the tumor boundary. A threshold density of tumor cell classifications is used to define the boundary (t-test; plotting individual cells from one mouse). **l-m.** Equivalent analyses in **j-k** with IBA1+ cells (microglia; n = 5 mice in **l**). \*\*\*\**p* < .0001.

To quantify the spatial distribution of S1PR3 and connect this to specific flow parameters, we stained GL261 tumor-bearing brains with our custom S1PR3 antibody and antibodies against a pan-astrocyte marker against ALDH1L1^23^ (Extended Fig. 3a) or IBA1 (Extended Fig. 3b). We trained a machine learning model using the open source software QuPath^24^ to classify S1PR3+ cells and cell types from immunofluorescence intensity and morphological features (Extended Fig. 3c). We also find that the percent of S1PR3+ astrocytes was negatively correlated with the distance to tumor boundary in all mice (Fig. 2j). Consistently, the distribution of S1PR3+ astrocytes is skewed toward the tumor boundary compared to S1PR3- astrocytes, demonstrated for one mouse (Fig. 2k). We also observe that the total number of astrocytes is negatively correlated with distance to the tumor border in all of the mice (Extended Fig. 3d). In contrast, while we find the percent of S1PR3+ microglia is negatively correlated with distance to the tumor boundary in all of the mice (Fig. 2l), this fails to result in a difference in the distribution of S1PR3+ and S1PR3- microglia across the tumor border (Fig. 2m). We do observe a negative correlation between the total number of microglia near the tumor border (Extended Fig. 3e). To incorporate flow, we superimposed these cellular coordinates with tumor-originating flow pathlines generated from the matched MRI data as in our previous work^19^. The percent of S1PR3+ astrocytes and the count of total astrocytes is significantly correlated with tumor originating pathline density in three of four mice, with a non-significant positive correlation for the fourth mouse (Fig. 2j; Extended Fig. 3d). In contrast, correlations are inconsistent between the percent S1RP3+ of IBA1+ cells or the total count of IBA1+ cells and tumor originating pathline density (Fig. 2l; Extended Fig. 3e). Thus, we hypothesized that there was a role for S1PR3 in astrocytes in peritumoral regions of flow, and, though we did not find consistent relationships between S1PR3+ microglia or the total count of microglia and flow, we did find them in higher numbers in the tumor-adjacent region where they would be expected to influence the microenvironment.

### Glial cells upregulate S1PR3 in response to fluid flow and shear stress in a rate-dependent manner

Since flow predicts S1PR3 positivity in parenchymal cells, we sought to identify whether experimental induction of flow induces S1PR3 reporter expression in naïve mice. We developed a method to infuse fluid into the mouse brain and track S1PR3-mCherry expression by intravital imaging (Fig. 3a). We infused fluid at defined rates, as confirmed by tracking of dextran-fluorescent tracers (Extended Fig. 4), and compared to the baseline signal before flow (Fig. 3b-c). Infusing at 1 µL/min induces a striking increase in mCherry signal that lasts at least nine days after the one-time infusion (Fig. 3d). Interestingly, this effect is not observed when infusing at 0.1 µL/min (to not induce pressure driven flow) or a higher 3 µL/min. To specifically correlate glial S1PR3 expression with flow, we disrupted the blood-brain barrier with focused ultrasound to increase fluid flow^25^. We then performed contrast-enhanced DCE-MRI to calculate flow magnitude (Fig. 3e; Extended Fig. 5a-e), harvested brains for immunofluorescence and classified cell types (Extended Fig. 5f-g), and superimposed the spatial features as before. Because of anatomical differences in cellular composition leading to noise, we segmented the isocortex for analysis (Extended Fig. 5f-g). Flow magnitudes reached approximately 1.2 μm/s, within the range in our tumor models, and the percent of S1PR3^+^ astrocytes significantly positively correlates with flow magnitude (Fig. 3f-h; Extended Fig. 5e).

**Figure 3.**
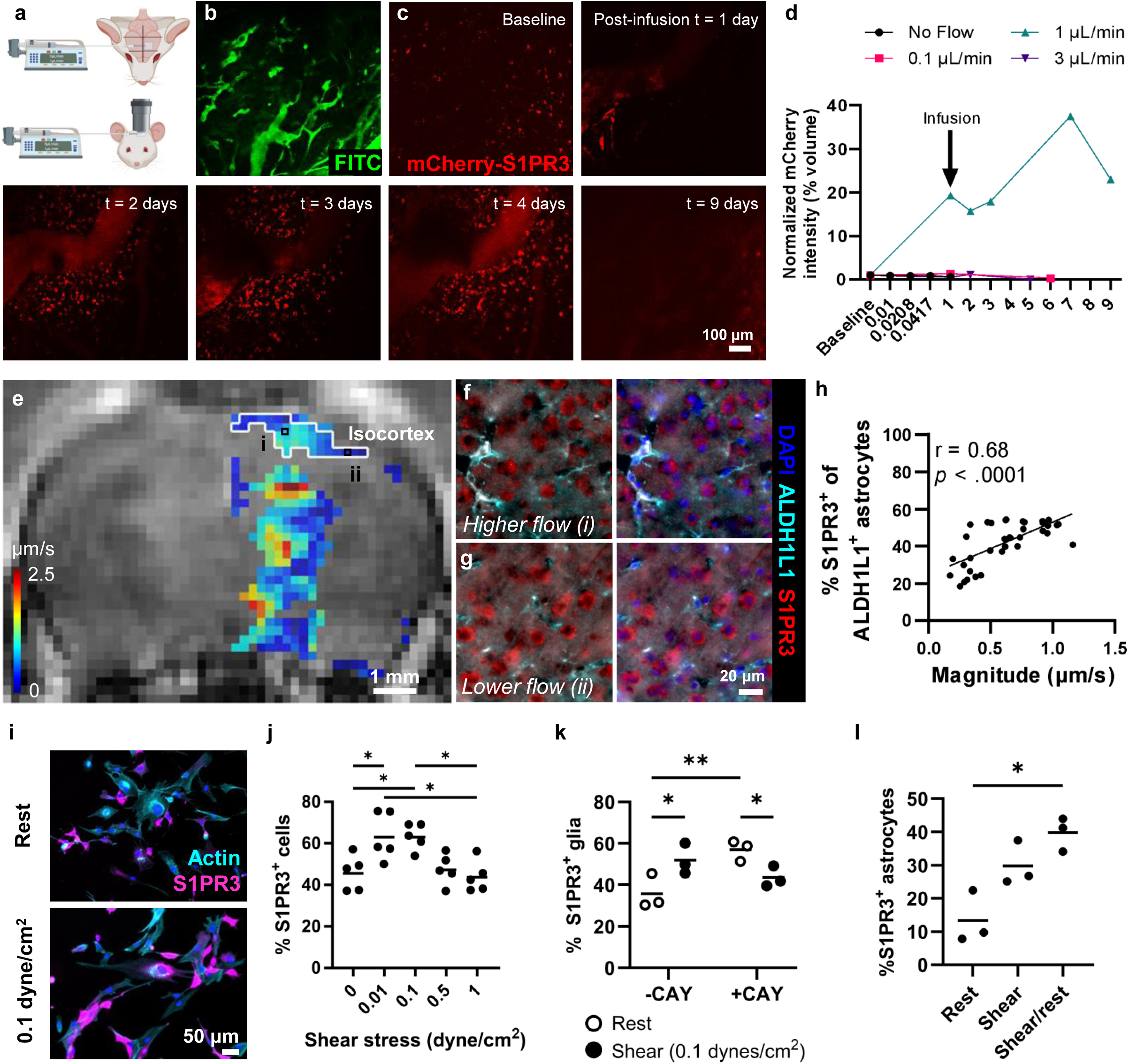
Glial cells elevate S1PR3 in response to enhanced fluid flow in tumor-naïve models. **a.** Experimental setup for live imaging through optical window of tumor-naïve S1P3^mCherry^ mice infused through an intracranial catheter into the parenchyma. **b.** Vascular labeling and **c.** reporter fluorescence during infusion. **d.** Threshold reporter fluorescence reported as S1PR3+ volume relative to the ROI volume and demonstrated for multiple pump infusion rates (n = 1-3 mice/group) **e.** Focused ultrasound-mediated blood-brain barrier opening paired with DCE-MRI (day 0) to calculate interstitial flow velocity in the contrast-enhanced region. **f-g.** Immunofluorescence staining (ALDH1L1 astrocytes) on brains harvested on day 2 post-sonication. **h.** Correlation between classified astrocytes expressing S1PR3 with flow (plotting MRI pixel values in isocortex shown in **e**; Pearson r) in a single mouse. **i.** Astrocyte and microglia co-culture in microfluidic model of laminar fluid shear stress (1 h) or rest control showing post-shear immunofluorescence. **j.** Quantified S1PR3+ cells relative to total glia across differing calculated shear stresses (1 h shear; n = 5 independent experiments; RM one-way ANOVA; Tukey’s multiple comparisons). **k.** Laminar shear on co-cultured glia (1 h at 0.1 dynes/cm^2^) showing effect of S1PR inhibition with CAY10444 (CAY) (n = 3 independent experiments; RM two-way ANOVA). **l.** S1PR3 expression in sheared astrocytes (1 h at 0.1 dynes/cm^2^) followed by overnight rest (n = 3 independent experiments; RM one-way ANOVA). *p < .05; **p< .01.

We used a 2D microfluidic model to evaluate the dependence of S1PR3 protein expression in human astrocytes and microglia on fluid shear stress magnitude, which is the surface force exerted by interstitial flow. Physiological flow in the brain is estimated to exert 0.01 dynes/cm^2^ of shear stress, whereas pathological flow is predicted to exert 0.6-2 dynes/cm^2^ ^10,26–28^. We find that human glial co-cultures exhibit a biphasic response to shear, with lowest S1PR3 immunoreactivity under static (no shear) and higher shear stress conditions (0.5-1 dynes/cm^2^) and a peak in S1PR3 at low pathological shear stress (0.1 dynes/cm^2^) (Fig. 3i-j). This increase in glial S1PR3 after 1 hour of 0.1 dynes/cm^2^ shear at least partially depends on S1PR3 signaling, since blocking the receptor using a small molecule significantly lowers the number of S1PR3+ glial cells (Fig. 3k). However, adding the inhibitor itself also increases baseline S1PR3 immunoreactivity. Interestingly, while astrocytes show a small increase in expression after 1 hour of shear, S1PR3 immunoreactivity continues to rise after an overnight rest period without shear (Fig. 3l). These data, in combination with our *in vivo* imaging results, suggest flow-stimulated S1PR3 expression in glial cells may involve receptor-mediated signaling that is sustained after the stimulus is removed. Ultimately, both our *in vitro* and *in vivo* results show glial cells do increase expression of S1PR3 in response to higher flow, but this response is not binary and depends on the flow rate and/or amount of shear stress.

### Glial S1PR3 regulates flow-enhanced glioma invasion

We recently published that interstitial flow correlates with invading tumor cells. Because we found interstitial flow at the tumor border also correlates with S1PR3+ expression in glia, particularly astrocytes, we hypothesized that glial expression of S1PR3 may correlate with and possibly influence tumor invasion. We find that S1PR3-mCherry signal intensity is significantly higher in peritumoral regions containing invaded GFP-GL261 compared to regions without invasion (Fig. 4a-b). To further understand the dynamics, we implanted GFP-GL261 tumors into S1PR3-mCherry mice and tracked the relationship between tumor progression and S1PR3 reporter expression by intravital imaging. mCherry expression is visibly enhanced surrounding the tumor at both 10 and 16 days after implantation (Fig. 4c). These results suggest there is a relationship between S1PR3 expression in parenchymal cells and tumor cell invasion.

**Figure 4.**
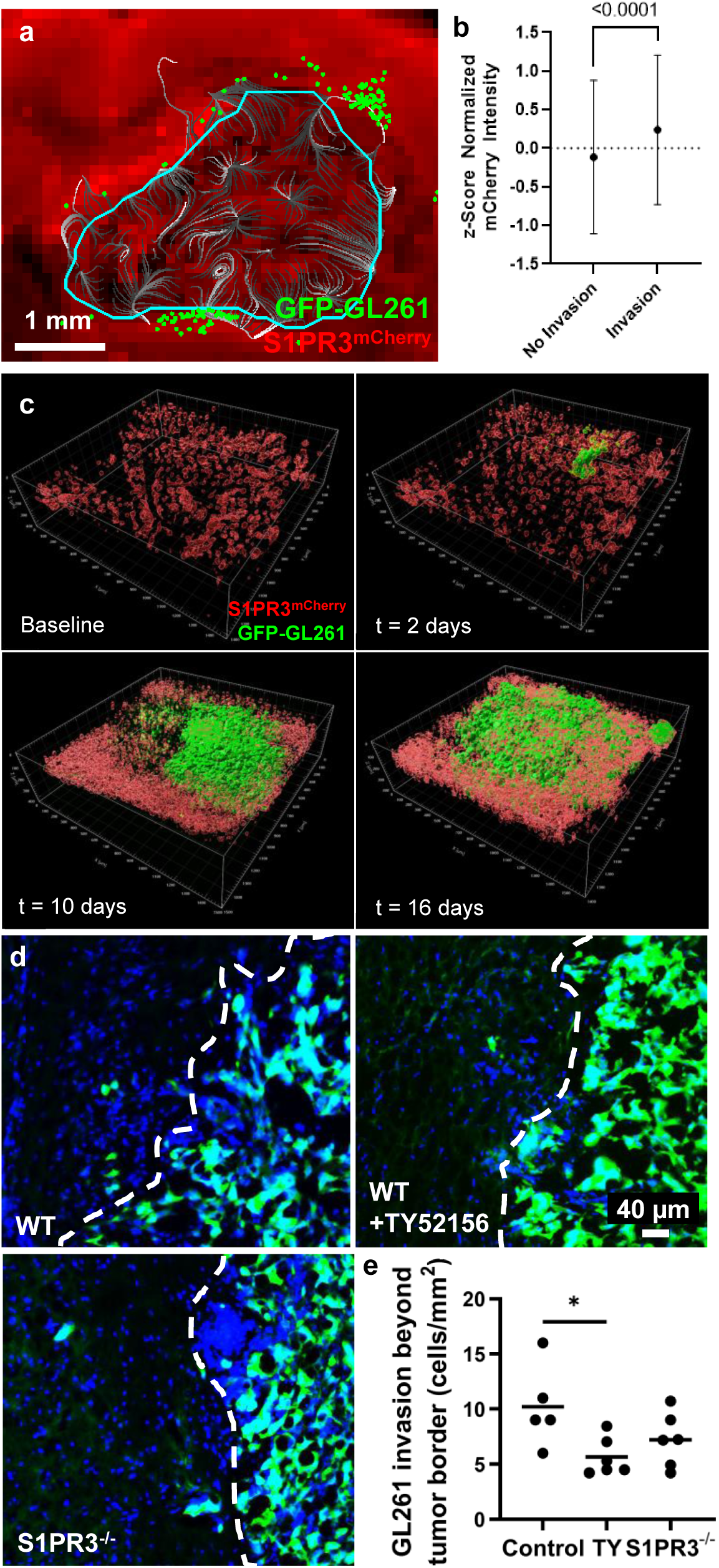
Peritumoral S1PR3 expression correlates with invasion and tumor progression, and targeting reduces murine glioma invasion. **a.** Intracranial implant of GFP-GL261 in S1PR3 reporter (S1PR3^mCherry^) mouse at day 15 post-implant, showing classified tumor cells invaded beyond tumor border (cyan) on a composite of three registered tissue sections. **b.** Reporter fluorescence intensity of MRI pixels containing at least one invading cell (invasion) or no invading cells (no invasion). **c.** GFP-GL261 bearing S1PR3^mCherry^ mice fitted with an optical window for progressive imaging of reporter fluorescence over time. **d.** GFP-GL261 implanted mice (C57Bl/6 or S1PR3^-/-^) under enhanced flow via convection enhanced delivery (day 7 post-implant) were treated every other day post-implant with 5 mg/kg intraperitoneal TY52156. Tissue was harvested on day 10 for sectioning and microscopy, showing tumor boundary (dotted line). **e.** Quantified (n = 5-6 mice/group; one-way ANOVA). \**p* < .05.

To further assess this relationship and test the therapeutic implications, we evaluated the effect of S1PR3 inhibition on tumor cell invasion in the syngeneic GL261 glioma model. We again used a catheter-based approach to exogenously enhance flow in the tumor, as previously described^29^, at a time when tumors were small enough to have minimal pressure gradient of their own (day 7). We started daily systemic administration of either the S1PR3-specific small molecule inhibitor TY52156^30^ or a vehicle control two days prior to adding flow. Three days after adding flow, we isolated the tissue and quantified tumor cell invasion by IHC. We find that blocking S1PR3 signaling with TY52156 in wild-type mice significantly reduces tumor cell invasion in the presence of increased convective flow, whereas knocking out in the microenvironment only (S1PR3^-/-^) did not confer a significant reduction in invasion (Fig. 4d-e). This dichotomy suggests there may be a benefit to targeting S1PR3 both in the microenvironment and on the tumor cells using a broadly-distributed small molecule inhibitor as compared to only targeting the TME.

We next assessed individual contributions of the tumor, glial cells, and flow on the effects of S1PR3 inhibition. To answer this question, we leveraged our tissue engineered model of the invasive glioma interface^5,17^(Fig. 5a). This tri-culture model uses a tissue culture insert, where invasion can be easily measured, and comprises human microglia, astrocytes, and patient-derived glioma cells in a 3D matrix under gravity-driven interstitial flow. For the G34 glioma stem cell line, the individual addition of interstitial flow, glial cells, or the S1PR3 inhibitor TY52156 did not affect glioma invasion, except in conditions involving covariates. Instead, invasion is inhibited only when the S1PR3 inhibitor is added in the presence of glial cells and interstitial flow (Fig. 5b). The effect of flow, glia, and S1PR3 inhibition is heterogeneous across multiple patient-derived glioma stem cell lines and the U251 glioma line, but the majority of cell lines we tested show reduced invasion under conditions with glia, flow, and the inhibitor (Fig. 5c; Extended Fig. 6a-c). Therefore, glial cells and flow mediate the effect of S1PR3 inhibition on invasion of glioma cells, but these responses may be patient-specific.

**Figure 5.**
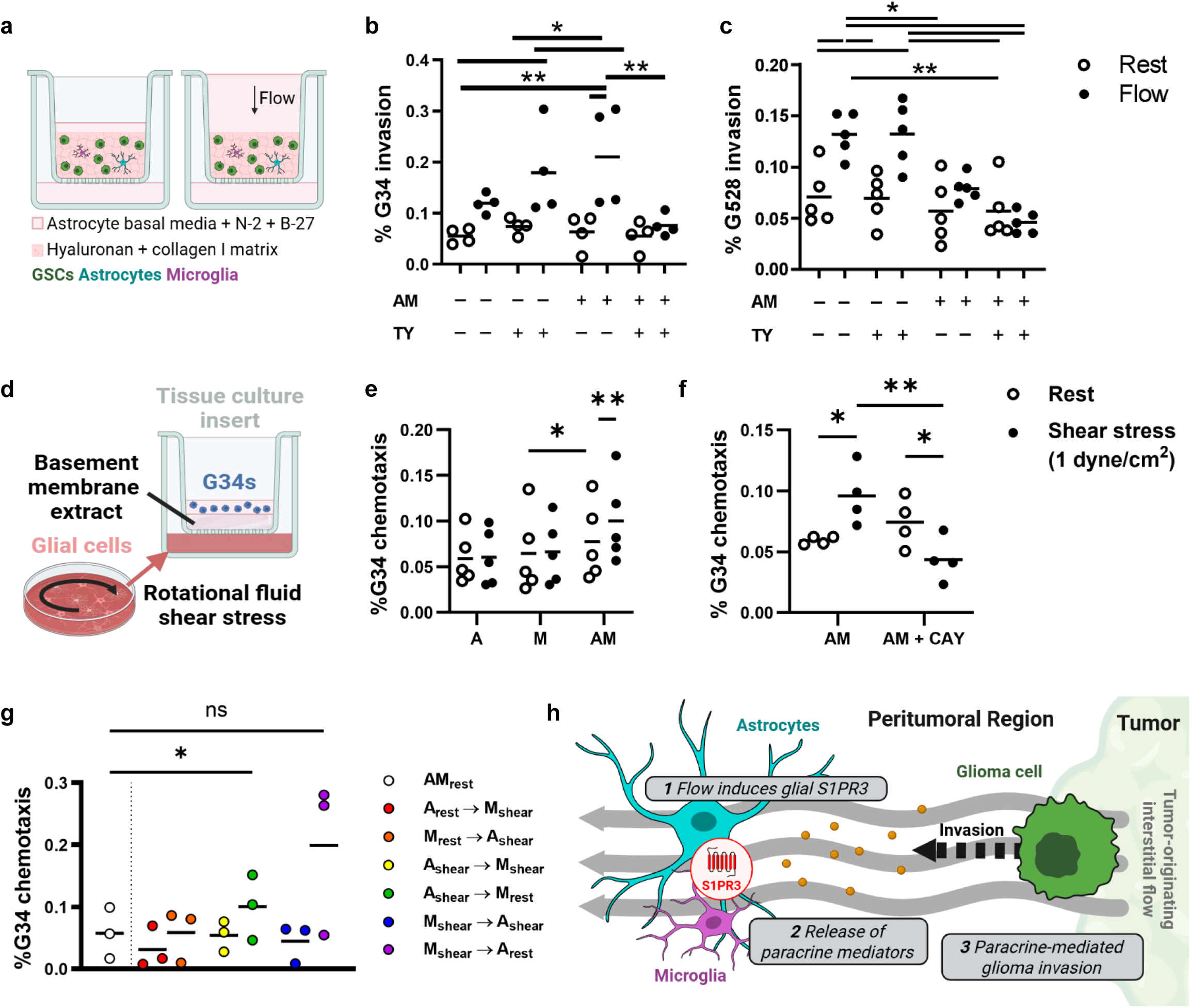
Glia regulate flow-enhanced glioma invasion via S1PR3. **a.** Patient TME model containing GSCs, astrocytes, and microglia in a 1.2 mg/mL collagen I, 2.0 mg/mL hyaluronan matrix in a tissue culture insert. A fluid pressure head (media) drives gravitational flow through the matrix. GSCs are permitted to invade through the porous membrane for 18-20 h following addition of the pressure head. **b-c.** Effect of flow, glia, and TY52156 treatment (10 μM) on G34 (**b;** n = 4 independent experiments) and G528 (**c;** n = 5 independent experiments) invasion in the TME model (one-way ANOVA with Tukey’s multiple comparisons). **d.** Chemotaxis model where mono- or co-cultured glia are sheared for 18-20 h at 1 dyne/cm^2^, then conditioned media is added to a well plate below a tissue culture insert containing GSCs suspended. in the hyaluronan-collagen gel. GSCs are allowed to invade through the porous membrane toward conditioned media for 18 h. **e.** Chemotaxis model in (**d**) with G34 exposed to shear-conditioned media from isolated or combined glia (n = 5 independent experiments; RM two-way ANOVA; Tukey’s multiple comparisons). **f.** Chemotaxis model in (**d**) with sheared co-cultured glia exposed to vehicle or S1PR inhibitor CAY10444 (CAY) (n = 4 independent experiments; RM two-way ANOVA; Fisher’s LSD). **g**. Glia were cultured alone or together under rotational shear or rest (1 dyne/cm^2^). Conditioned media was transferred to the second cell type indicated by arrow, then used for the chemotaxis model in (**d**) (n = 3 independent experiments; RM one-way ANOVA; Dunnett’s multiple comparisons). **h**. Schematic of hypothesized mechanism where glial S1PR3 regulates glioma invasion during flow. A, astrocytes; M, microglia; \**p* < .05; \*\**p* < .01.

Towards determining if this effect is mediated by soluble factors, we collected conditioned media from human glial cells cultured alone or together in 2D under 1 dynes/cm^2^ of rotational shear and conducted chemotaxis assays with the patient-derived glioma stem cell lines (Fig. 5d). G34, but not G528, invades more toward sheared conditioned medium from co-cultured glial cells but not individual glial cells (Fig. 5e; Extended Fig. 6d). This effect also appears to be transient, since conditioned media collected the day after shearing is no longer chemotactic (Extended Fig. 6e). Importantly, treating glia with the S1PR3 antagonist CAY10444 during shear significantly reduces the resulting G34 chemotaxis (Fig. 5f). Both G2 and G528 were again not responsive to sheared glial cells, but CAY10444 treatment did reduce G2 invasion in response to sheared glia (Extended Fig. 6f-g). Fluid shear therefore induces astrocytes and microglia to secrete factors that influence the motility of some, but not all, patient glioma cells through a mechanism involving glial S1PR3. Shear-stimulation of astrocytes may be the initiating factor, since the chemotactic effect is only observed for conditioned media from sheared astrocytes that was transferred to microglia prior to running the chemotaxis assay (Fig. 5g). Collectively, our data support an inter-glial signaling mechanism that regulates flow-enhanced glioma invasion, whereby fluid shear stress on glial cells induces the release of unknown paracrine mediators that interact with glioma cells to influence invasion (Fig. 5h).

### S1PR3 expression relates to interstitial flow dynamics and glioma invasion in patients

To assess the potential prognostic value of S1PR3 expression in the clinic, we queried existing databases to define the basic characteristics of S1PR3 expression across GBM patients and analyzed whether these correlate with clinical outcomes. Using the GEIPA database, which pools RNAseq data from TCGA and GTEx^31^, we find S1PR3 expression is significantly higher in GBM tissue compared to non-diseased postmortem human brain tissue (Extended Fig. 7a). There is also a significant correlation between S1PR3 protein abundance and time to recurrence, and a weak trend with overall survival, when incorporating clinical outcomes from mass spectroscopy data collected by the Clinical Tumor Proteomic Analysis Consortium (CTPAC)^32^ and available via cBioPortal^33–35^ (Extended Fig. 7b-c). We classified patient samples into groups for S1PR3-high and S1PR3-low based on protein abundance z-scores ± 1. S1PR3-high samples exhibit significantly lower overall survival compared to S1PR3-low samples (Exteded Fig. 7d). There is also a trend toward faster recurrence in S1PR3-high versus -low samples (Extended Fig. 7e). Thus, S1PR3 expression in bulk tumor tissue may be a useful biomarker to predict disease progression. We find no significant interactions between various demographic features and S1PR3 across cohorts (Extended Fig. 7f-i).

We next compared S1PR3 immunostaining in tumor-adjacent normal tissue against pathologist-identified tumor infiltrative regions to understand whether S1PR3 expression is replicated in the invasive microenvironment from patient GBM samples. S1PR3 reactivity was clearly visible in the infiltrative region compared to normal tissue (Fig. 6a). We analyzed 34 patient tumor infiltrative regions and corresponding survival data using a custom antibody against the amino-terminus of human/mouse S1PR3 (Extended Fig. 7j). Twenty-one of the sample cores were S1PR3-positive and thirteen were S1PR3-negative (Extended Fig. 7k). Interestingly, we also identify S1PR3 staining in breast and melanoma brain metastases (Extended Fig. 8). These data suggest that though there are S1PR3+ regions peritumorally, there is heterogeneity across tissue samples and could depend on the status of fluid flow in the biopsied region.

**Figure 6.**
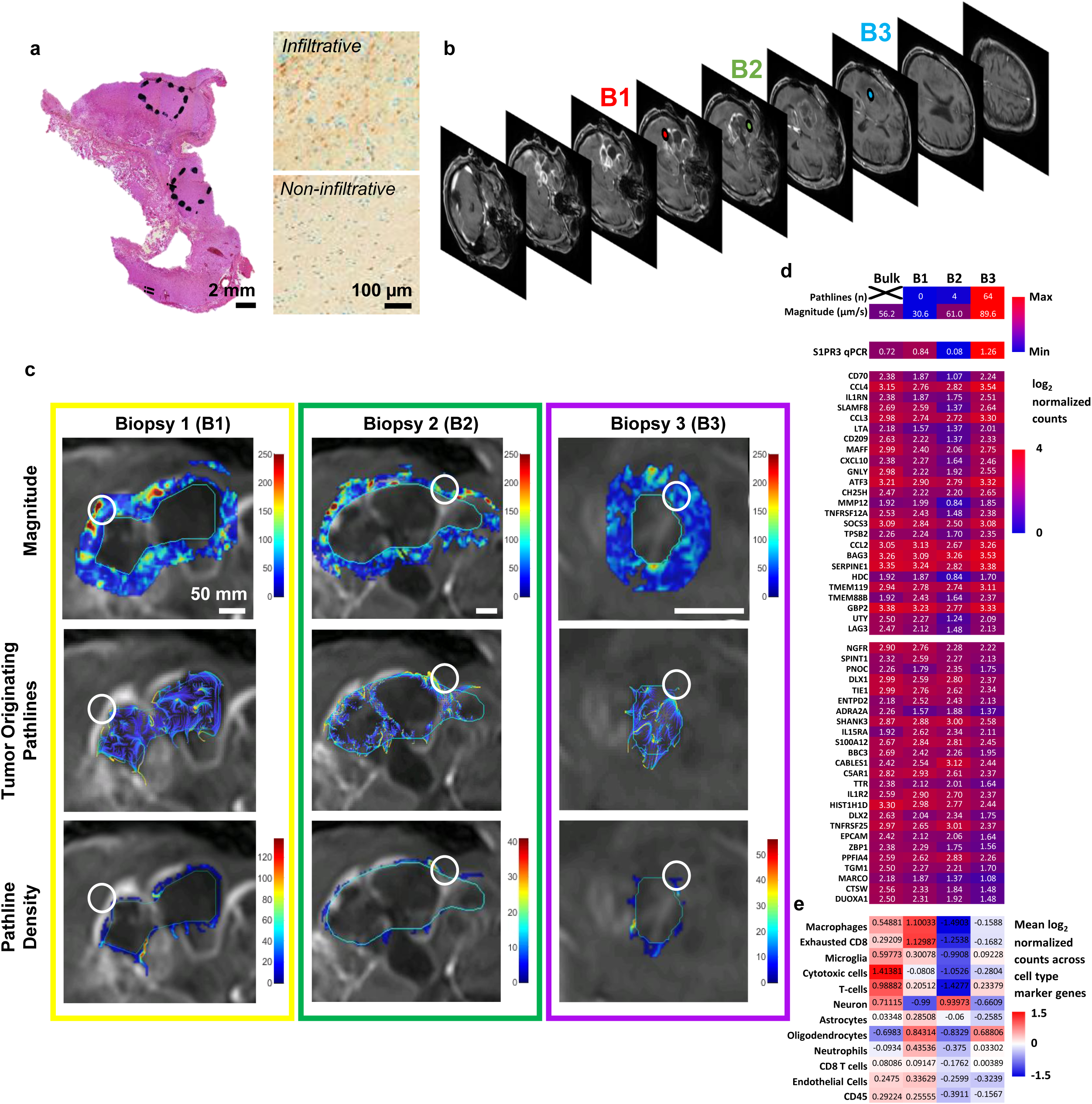
Image-localized biopsies from a patient with GBM are consistent with enhanced S1PR3 in high-flow peritumoral regions. **a.** H&E and DAB staining for S1PR3 in a tumor-infiltrating region vs. normal tissue in a patient tumor. **b.** Stacked T1 image from patient with diagnosed GBM indicating MRI-guided biopsy locations (B1-3). **c.** DCE-MRI quantified interstitial transport metrics in the tumor-adjacent region superimposed on T1 image for each biopsy location. Pathlines are calculated from the interstitial fluid velocity vector field and indicate flow trajectories from the tumor to parenchyma. **d.** Heatmap combining interstitial transport metrics, RT-qPCR for S1PR3, and neuroinflammation gene expression microarray across each biopsy. Top gene and cell hits are sorted by the ratio of B3 (highest flow) over the geometric mean of B1 and B2. **e.** Rosalind analysis for individual cell types based on the neuroinflammation panel to identify different cell types in the individual biopsy.

Determining if peritumoral S1PR3 expression in patients is significantly linked to invasion in flow regions was not possible using current databases or prior patient biopsies because either the tissue location is within the bulk or not anatomically known relative to the MRI. In addition, many MRI protocols do not allow for flow analysis^36^. Thus we performed MRI conducive with applying flow analysis prior to a resective surgery. During surgery, we performed MRI-localized biopsies, collecting three peri-tumoral biopsies (B1-3) and one bulk tumor sample (Fig. 6b-c). We quantified S1PR3 expression by RT-qPCR and immunological features using the nCounter neuroinflammation panel (Fig. 6d). Though our sample size is too limited for statistical analysis, the biopsy with the highest S1PR3 expression (B3) also exhibited the highest flow magnitude and highest number of tumor-originating pathlines compared to the other two peritumoral and tumor bulk samples. Both B3 and B2 appear to contain a lower representation of peripheral immune cells compared to B1 and surprisingly, a reduced astrocyte signature (Fig. 6d; Extended Fig. 9). While a larger study is needed, these preliminary data suggest there is indeed heterogeneity in S1PR3 expression in the peritumoral space, with S1PR3 expression being increased in peritumoral regions of flow in human GBM.

## Discussion

The phenotypic plasticity of glioblastoma cells in response to diverse microenvironmental cues promotes an adaptable, invasive, and difficult to target cancer with limited treatment options. Previous work has tied glioma invasion to broad patterns in interstitial fluid flow across the brain^7^ and peritumoral flow^8,19^. The capacity of flow to influence glioma invasion and motility has been demonstrated by multiple groups^8,10,14^, complementing findings across cancer types^37^. However, an open area of research is how this flow affects brain parenchymal cells and whether that also influences glioma invasion. In this paper, we sought to consider the impact of the TME in mechanisms driving flow-enhanced glioma invasion and identify candidate therapeutic targets to limit this effect. We identified an intercellular mechanism whereby flow-stimulation of astrocytes and microglia, marked by expression of sphingosine-1-phosphate-receptor 3 (S1PR3), induces glioma motility.

Our study is not the first to connect S1PRs and flow. In fact, one work implicated S1PRs in fluid mechanosensing^21^. Jung et al. 2012 identified a critical role for S1PR1 in regulating vascularization and blood vessel permeability. Fluid shear stress induced assembly of adherens junctions and could be inhibited with FTY720-P, the bioactive form of fingolimod, an FDA-approved S1PR modulator. Though our report is the first involving S1PR3 in flow-induced invasion and migration, S1PR3 has been implicated in invasion and migration across cellular contexts. Abundant literature exists implicating S1PR3 in immune cell migration, tied to the dependence of many cell types on systemic sphingosine-1-phosphate (S1P) gradients for lymphatic homing, reviewed in^38^. Emerging literature has also tied S1PR3 to cancer invasion within the brain. An initial *in vitro* study by Young and Van Brocklyn, 2007^39^ implicated multiple S1PRs, including S1PR3, in glioblastoma cell invasion through Matrigel. Recently, astrocytic S1PR3 was shown to regulate blood-tumor-barrier permeability and metastasis of murine breast tumors to the brain^40^. This effect was also attributed to secretion of paracrine mediators. An influx of fluid from an enhanced blood-brain barrier would be expected to influence interstitial flow composition (soluble factors and viscosity) and velocity. That this is an S1PR3+ astrocyte-mediated mechanism is particularly interesting and could reflect a congruent intercellular mechanism to our findings whereby flow on glial cells, especially astrocytes, induces secretion of a paracrine factor that increases glioma chemokinesis, dependent upon glial S1PR3 signaling.

Increased parenchymal S1PR3 expression by flow in tumor-naïve brains and tissue cultures suggests a context-agnostic mechanism applicable across neuropathologies. This possibility is consistent with the aforementioned report of astrocytic S1PR3 mediating blood-brain-barrier permeability in breast-to-brain metastases^40^ and the efficacy of S1PR antagonism (CAY10444) in ischemia-reperfusion injury related to acute S1PR3 upregulation in perivascular astrocytes^41^. Further, S1PRs are known immunological modulators, originating from work with fingolimod in multiple sclerosis^42,43^. Though fingolimod’s immunosuppressive effect was initially attributed to lymphopenia, or blunting lymphocyte egress from the lymph node^44^, emerging work across neuropathologies has highlighted additional effects of fingolimod on neuroinflammation in connection to astrocytes^45–48^, including via S1PR3^48,49^. A particularly relevant study demonstrated that fingolimod treatment blunted the response of astrocytes to fluid shear stress in the presence of sphingosine-1-phosphate^50^. In a similar vein, S1PR3^-/-^ or S1PR targeting improved neuroinflammatory phenotypes in the ischemia-reperfusion study discussed previously, including a non-significant but reduced mean albumin accumulation, an indicator of vascular leakage^41^. Besides S1PR3-regulated responses, fluid shear stress was recently shown to induce a neurotoxic phenotype in astrocytes and microglia via purinergic signaling^51^. The extent to which these pathways may or may not overlap and represent a shared, innate astrocytic mechanism that regulates interstitial flow and neuroinflammation is not yet understood, but it is an intriguing area of further research with translational potential across pathologies. Notably, however, the direct effect of S1PR modulation on astrocytes must be carefully uncoupled from the inflammatory effects of, and crosstalk with, infiltrating peripheral cells^2^.

An important feature of our approach is that it demonstrates the necessity/utility of integrating biophysical forces into tissue engineered models for drug discovery and development. S1PR3 inhibition was only effective for reducing invasion in the presence of both elevated fluid flow and glial cells. This indicates that the response of the glial cells to the mechanical deformations exerted by fluid shear activated mechanisms in the glioma cells that could not be modeled in a monoculture or static system. Using this same model, we previously showed that fluid flow is a major modulator of glioma proliferation and expression of stem-like markers^5^. These complimentary findings serve as an initial validation for the relevance of this model system, and we hope this system and others like it may facilitate drug discovery pipelines and understanding complex intercellular mechanisms.

To gauge the potential clinical applicability of S1PR3 targeting, we collected image-localized biopsies and compared gene expression arrays and S1PR3 qPCR to patterns of peritumoral flow. The biopsy with the greatest tumor-originating flow did indeed express the most S1PR3. Our previous work indicates that flow predicts disease progression in animal models^19^, and tracking flow in patients is a minor addition to the current standard of care imaging protocol with potential major implications. Further work to stratify patients likely to be sensitive to pharmaceutical targeting of flow, e.g., against S1PR3, is a promising area of future study. Given the heterogeneous sensitivity to blocking this pathways across patient cell lines, it may be beneficial to combine with blockade of other receptors and ligands that promote invasion in response to flow, such as CXCR4/CXCL12 and CD-44^14^. In literature outside of glioma, fluid shear stress and flow-directed soluble factors are known to influence a broad array of other signaling pathways, such as ECM-actin coupling via integrins and mechanosensitive ion channels, junctional proteins, and G-protein coupled receptors (besides S1PR3)^53–57^. These mechanisms still need to be investigated in the context of enhanced peritumoral flow in GBM. Of particular interest would be other S1PRs; S1PR1 and S1PR3 are sensitive to fluid shear stress^21,58^, but whether the other S1PRs are mechanosensitive is unknown. To this end, a targeted expansion of our MRI-matched patient biopsy data would be highly informative for identifying additional therapeutic candidates for personalized therapy.

Most interesting across all of our models and systems is the spatial and cellular distribution of S1PR3 expression and activity. For example, we observed a range of sensitivity to S1PR3 inhibtion across patient-derived cell lines. In mouse models, there is a correlation between S1PR3 with flow, but flow is inherently spatially distributed and dynamic. Hence S1PR3 inhibition may specifically work in peritumoral regions of high flow or may need to be targeted to these regions for maximal effect. Further, in a single patient, we similarly see intra-peritumoral heterogeneity of S1PR3 expression based on a series of biopsy samples. These do appear to correlate with flow, similar to our mouse studies, but further evaluation of these correlations and their impacts are needed to truly understand the benefits and limitations of S1PR3 inhibition in patients In spite of these caveats, the role of S1PR3 in invasion of glioma and its specific activity in flow regions presents a unique interaction between the microenvironment and glioma cells and a potential target for future development.

## Methods

### Cell culture

Patient-derived human glioblastoma stem cells (GSCs) were a generous gift to Dr. Benjamin Purow from Dr. Jakub Godlewski and Dr. Ichiro Nakano, who derived them while at Ohio State University^20^. These cells (G2, G34, G62, and G528) were maintained in non-treated culture flasks in Neurobasal medium (Life Technologies) supplemented with 1% B27, 0.5% N2, 0.01% FGF, 0.1% EGF, 0.3% L-Glutamine, and 1% penicillin-streptomycin. Human primary cortical astrocytes (Sciencell) were cultured according to the manufacturer’s suggested protocol. Human SV40-immortalized microglia (Applied Biological Materials, Inc.) were cultured in Dulbecco’s Modified Eagle’s Medium (DMEM; Life technologies) supplemented with 10% fetal bovine serum. Both astrocytes and microglia were seeded on tissue-culture treated flasks coated with rat-tail collagen I (Corning 354236); approximately 3 uL/cm^2^ collagen was evenly spread across the growing area and incubated for 30 minutes at 37°C followed by a wash in PBS 5 minutes at 37°C. The murine glioma cell line GL261 was previously transduced to express green fluorescent protein (GFP) and periodically purified by selection with 2 μg/mL puromycin (Thermo Fisher)^29^. GFP-GL261 were maintained in tissue-culture treated flasks in DMEM + 10% FBS as described above. All cell lines were maintained at 37°C in a humidified incubator containing 5% CO_2_ and 21% O_2_. Cell lines were tested annually for mycoplasma.

### Intracranial tumor implantation

Tumor implantation protocols were approved by the Institutional Animal Care and Use committees at University of Virginia (protocol 4021) and Virginia Polytechnic Institute and State University (protocols 17-113, 20-146, and 20-183). Implanted mouse strains include Non-obese Diabetic Severe Combined Immune Deficiency (NOD SCID) (NCI), C57BL/6 (The Jackson Laboratory), B6.Cg-S1pr3tm1.1Hrose/J (S1P_3_ ^mCherry^, The Jackson Laboratory 028624; C57BL/6 background)^22^, and S1PR3^-/-^ (C57BL/6 background, a generous gift from Dr. Mark Okusa)^59^. Patient-derived xenografts were established in male NOD SCID mice using either 10,000 G2, 10,000 G34, or 400,000 G528. Syngeneic tumors were established in C57BL/6, S1PR3^-/-^, or S1P_3_ ^mCherry^ mice using 100,000 GFP-GL261.

8-10 week old mice were prepared for aseptic surgery and secured on a stereotactic frame (Steolting). For the glial S1PR3 localization vs. flow study in GFP-GL261 implanted S1P_3_^mCherry^ mice, 8-11 month old mice were used. The skull was exposed using a small incision, and a burr hole was drilled in the skull at 2 mm lateral and posterior to bregma for tumor inoculation. Tumor cells were stereotactically injected at a depth of 2.2 mm at 1 uL/min for 5 mins. After inoculation, the burr hole was filled with dental wax, the incision sutured, and the animal was allowed to recover in the home cage. Analgesics are administered prior to surgery and for 48 hours after surgery for pain management. For the S1PR3 inhibition study, TY52156 was resuspended in DMSO at 5 mg/mL, and drug or vehicle control was administered intraperitoneally (5 mg/kg) every other day starting the day after tumor implantation until study termination. The survival study in S1PR3^-/-^ mice was terminated by CO_2_ asphyxiation following humane endpoints including > 10% loss of body mass.

### Murine patient-derived xenograft tissue harvest for microarray

Murine patient-derived xenografts from the GSC lines G2, G34, and G528 were given tail vein injections of 10 mg/mL (5 mg/kg) Evans blue the day prior to euthanasia following our previous studies^18^. Evans blue intensity correlates with regions of outward flow^18^. Following euthanasia, brain tissue was collected and transported to the imaging suite in cold RNA*later™* (Invitrogen) and cut in half coronally at the injection site. Approximate areas of ‘high’ and ‘low’ interstitial flow out of the tumor were located by comparing prior magnetic resonance images with Evans blue dye visualized in brightfield and RFP channels on an AxioZoom stereoscope (Zeiss). Using this collective information, regions of high flow, low flow, normal or control tissue, and tumor tissue were microdissected using fine forceps and stored frozen in RNAlater. The RNA is isolated using the PureLink™ RNA mini kit (Invitrogen) and submitted for microarray at the Bioinformatics Core at the University of Virginia. Microarray was performed by the UVA DNA Sciences core using the Affymetrix GeneChip Array (Mouse Gene 2.1 ST Array). Gene lists were generated from microarray data corresponding to genes >1.25 LogFC and <0.75 logFC were used in downstream analyses.

### Small animal magnetic resonance imaging for interstitial transport metrics

For GFP-GL261 tumor-bearing mice, DCE-MRI was performed on mice with detectable tumors on post-implantation days 15 and 24^19^. Mice were maintained under anesthesia with isoflurane (3% for induction and 1.5% for maintenance) in a 50:50 mixture of air-O_2_. Mice were imaged with a 9.4 T small animal MRI (Bruker BioSpec AVANCE NEO 94/20 USR, Ettlingen, Germany) equipped with 660 mT/m high power gradient. An active detunable 86 mm volume coil was used as the transmit coil and a planar 20 mm receive-only surface coil was used as the receive coil. T2-weighted images were acquired to visualize anatomical features and/or confirm tumor growth in GFP-GL261 mice. For the GFP-GL261 mice, T2 parameters were as follows: repetition time (TR) = 1800 ms, echo time (TE) = 40 ms, field of view (FOV) = 19.2 x 19.2 mm with a 192 x 192 matrix, slice thickness = 0.5 mm, number of slices = 16 with 9 averages, total acquisition time = 6.5 min/mouse. For the focused ultrasound study, the T2 parameters were: TR = 2500 ms, TE = 33 ms, FOV = 51.2 x 51.2 mm with a 256 x 256 matrix, slice thickness = 1 mm, number of slices = 14 with 7 averages, total acquisition time = 9.5 min/mouse. A pre-contrast T1-weighted image was collected for baseline signal normalization. Then, gadolinium (Magnevist, Bayer HealthCare Pharmaceuticals) was delivered via the catheter at a concentration of 0.1 mmol/kg in sterile, heparinized saline. Multiple post-contrast T1-weighted images were acquired over time using a fast low angle shot (FLASH) sequence. For the GFP-GL261 mice, T1 parameters were: n post-contrast T1 = 4, TR = 180 ms, TE = 11 ms, number of slices = 16 with 7 averages, total acquisition time = 3 min/mouse. For the focused ultrasound study, T1 parameters were: n post-contrast T1 = 5, TR = 190 ms, TE = 4 ms, number of slices = 14 with 5 averages, total acquisition time = 3 min/mouse. FOV, matrix size, and slice thickness were equivalent to T1 images. For patient-derived xenografts, DCE-MRI was performed following previously published protocols^14,18^. Interstitial transport metrics were quantified in MATLAB using the open-source tool Lymph4D following previous protocols^19,60^.

### Murine tissue harvest and sectioning

Mice were near-asphyxiated with CO_2_ and perfused intracardially with ice-cold saline or phosphate buffered saline. Mice for immunohistochemistry were subsequently perfused with 4% paraformaldehyde to fix the tissue and brains were dissected. After fixation, brains were post-fixed in 4% paraformaldehyde overnight at 4°C and cryopreserved in 30% sucrose. Tissue was frozen en bloc in Optimum Cutting Temperature (OCT) medium at -80°C and sectioned at 12 um on a cryostat (Leica Biosystems, Wetzlar, Germany). When relevant, tissue sections were matched to T2-weighted MRI based on anatomical features (i.e., dentate gyrus, white matter, ventricles, and tumor shape when relevant).

### Custom α-S1PR3 antibodies

To improve S1PR3 staining quality, we generated two custom antibodies against S1PR3, α-human/mouse S1PR3 antibody against a C-terminal peptide (CZ SRS KSS SSN NSS HSP KVK), and α-human S1PR3 against N-terminal residues 3-27 (RG NET LRE HYQ YVG KZC). Antibodies were commissioned through Aves Labs (Davis, CA, USA). Hens were injected with KLH conjugates of the peptide, and eggs were collected. IgY fractions were isolated and affinity purified.

### Immunofluorescence in murine tissue

For all studies, tissue was thawed from -80 °C, outlined with a hydrophobic PAP pen, serum blocked for one hour at room temperature, and incubated in primary antibodies in blocking solution. Primaries were washed 3X in PBS or PBS + TX-100, then an appropriate secondary antibody was added for one hour at room temperature (Alexa Fluor Life Technologies, Invitrogen, Jackson ImmunoResearch Laboratories, Inc., either Alexa Fluor or Dylight). Tissue was washed multiple times, and nuclei were labeled with DAPI. Coverslips were mounted with Fluoromount G (Southern Biotech, USA) on 1.5 mm coverglass. Specific protocols are described below.

#### mCherry cellular localization study in S1P ^mCherry^ mice bearing GFP-GL261 tumors

Tissues were blocked in 3% serum from the host of the secondary antibody. Primary antibodies were applied overnight at 4°C and included: α-mCherry (Abcam ab205402), α-GFAP (Abcam ab7260), α-Iba1 (Abcam ab5076), α-Oligodendrocyte specific protein (Abcam ab53041), α-CD31 (Abcam ab23864), α-PDGFR-β (Abcam ab32570), α-GLT-1 (Abcam ab106289). Slides were imaged using a 40x oil immersion objective on the LSM 800 confocal microscope (Zeiss). Images were taken in areas surrounding the tumor with high mCherry expression. mCherry signal was enhanced with the Alexa Fluor™ 555 Tyramide SuperBoost™ Kit (Invitrogen B40923).

#### S1PR3 glial localization vs. flow study in GFP-GL261 mice

Slides were placed on a hot plate at the lowest setting (≈ 30 °C) for 15 minutes. Slides were boiled in 1X pH 6 Antigen Unmasking Solution, Citrate-Based (Vector Laboratories, Inc., USA) for 10 min in a 900 W microwave. Slides were cooled at room temperature for 1 hour, then washed in 1X PBS. Blocking solution contained 1% bovine serum albumin, 0.3% TX-100, and 2% donkey serum. Primary antibodies were applied overnight at 4°C and included: α-S1PR3 (custom C-terminal antibody, Aves, 5 µg/mL) α-GFP (Novus Biologicals NB100-62622, 50 µg/mL), then either α-ALDH1L1 (Sigma-Aldrich MABN495, 5 µg/mL) or α-IBA1 (Synaptic Systems 234-308, 2 µg/mL). Whole slides were imaged on a slide scanner (Olympus Slideview VS200, Evident, Tokyo, Japan).

#### Focused ultrasound study

Slides were placed on a hot plate at the lowest setting (≈ 30 °C) for 10 minutes. Slides were immersed in 1X pH 6 Antigen Unmasking Solution, Citrate-Based (Vector Laboratories, Inc., USA) in a coplin jar and placed in a boiling water bath for 30 min. Slides were cooled at room temperature for 1 hour, then washed in 1X PBS. Blocking solution contained 2% donkey serum and 0.03% TX-100 in PBS. Primary antibodies were applied at room temperature for 2 hours: chicken α-S1PR3 (custom C-terminal antibody, Aves, 1 µg/mL), α-ALDH1L1 (Sigma-Aldrich MABN495, 5 µg/mL), and α-IBA1 (Synaptic Systems 234-308, 2 µg/mL). Whole slides were imaged on a slide scanner (Olympus Slideview VS200, Evident, Tokyo, Japan).

#### GFP-GL261 invasion study

Blocking solution contained 3% serum from the host of the secondary antibody in 0.3% TX-100. GFP fluorescence was amplified by staining with α-GFP (Life Technologies A21311, 10 µg/mL). Sections were imaged at 20X magnification on an EVOS FL Auto.

### Quantitative spatial immunofluorescence

Cell types and S1PR3^+^ or S1PR3^-^ cells were defined using the open-source software QuPath versions 0.4.3 (GFP-GL261 study) or 0.5.1 (FUS study)^24^. Nuclei were segmented using the deep-learning algorithm Stardist with the DSB2018 heavy augment model^61^ and expanded at a defined radius approximating the cell border. Compartment-specific (nucleus, membrane, and whole cell) features were calculated for each stain, such as size and shape and fluorescence intensity (mean, max, standard deviation, etc.). Additionally, to account for variations in background tissue fluorescence and positive cell staining intensities, textural (Haralick^62^), smoothed, and intensities for regional expansions (i.e., 25-100 μm circles) were calculated. Appropriate features were selected to train a random forest models to classify cells as GFP-GL261 tumor cells, ALDH1L1^+^ astrocytes, IBA1^+^ microglia/macrophages, or S1PR3^+^ or S1PR3^-^ cells. To quantify the distance between classified cells and the tumor border, full-tumor annotations were created from a density map of all tumor-classified cells and filled to remove holes. This border was expanded 400 um to define the peritumoral region.

### Correlating spatial immunofluorescence with interstitial transport metrics

Classified cell coordinates and raw immunofluorescent stains were exported and transformed to match MRI coordinates. Control point-registration (local weighted mean geometric transformation) was used to align tissue sections and MRI slices in Matlab Version R2021b. At least 12 control points were selected, referencing broad anatomical structures (ventricles, longitudinal fissure, and the cortical surface). Immunofluorescence-based metrics were registered to the Lymph4D output using the resulting transformation matrix. Tumor boundaries were defined from composite tissue sections as described in^19^.

### Intravital imaging experiments

Intravital imaging experiments were conducted at Virginia Polytechnic Institute and State University under approved protocol #20-183. B6.Cg-S1pr3tm1.1Hrose/J mice (S1P_3_^mCherry^, The Jackson Laboratory #028624) underwent cranial window surgery followed by implantation of an external catheter (P1 Technologies, C315GS-4/SPC) through the contralateral hemisphere such that the catheter tip was situated underneath the window and accessible for infusion when under the microscope. Mice were anesthetized with isoflurane and secured to a stereotactic frame (Kopf, Model 900). The scalp was removed along the midline and above bregma to expose the skull. A drill (Foredom, K.1070 and bit, Roboz Surgical Instrument Company, RS-6282C-33-1/2) was used to cut an approximately 2 mm x 2 mm portion of the skull to expose the brain, with the window placed on the left and caudal side of bregma. A coverslip was placed on top of this opening and secured using tissue adhesive. A burr hole was then drilled (with bit - Fine Science Tools, 19008-14) on the contralateral side approximately 2.3 mm away from the right edge of the window and a cannula placed into this opening and positioned underneath the cranial window using the stereotactic arm. Finally, everything was secured in place with dental cement and the animal allowed to recover for four days before imaging is performed. The mice were then imaged under isoflurane using either an Olympus FV1000 MPE with a XLPLN25X/1.05 NA water-immersion objective (Olympus) and band-pass emission filters: 630/92 and 520/70 (Semrock) or a Nikon A1R multiphoton microscope with a 25X/1.10 N.A. W water-immersion objective (Nikon), a tunable femtosecond pulsed Ti:Sapphire laser (Coherent MRU X1), and four band-pass emission filters: 440/80, 525/50, 641/75, and 732/68 (Semrock). Mice were secured to a 3-axis stereotactic frame (Kopf, Model 1900) during imaging.

#### Infusion study

Seven mice (1-3 per group) were injected retro-orbitally with a fluorescent dye, FITC dextran, to label vasculature and ensure imaging of the same area during each imaging session. For infusion, a syringe pump (Kent Scientific, GenieTouch) was used to inject FITC dextran through a cannula (P1 Technologies, C315IS-4/SPC) that was inserted into the surgical catheter. The flow rate of the syringe pump was varied between 0.1 μL/min, 1 μL/min, and 3 μL/min and a total volume of approximately 5 μL delivered. Additionally, one group of mice received no infusion. Imaging was performed with the same laser settings for each session per mouse. Time lapse images were taken during each infusion of a location near the catheter to visualize dextran entering and moving through the tissue. Overview images of the whole window area were taken as a tile scan before and after infusion and follow up images of tile z-scans were also acquired at various time points post-infusion. Analysis was performed using Imaris software to quantify the mCherry signal at each time point.

#### Tumor progression study

Four days after cranial window surgery, 200,000 GFP-GL261 cells (100,000 cells/μL) were infused through the catheter at a rate of 0.333 μL/min and total volume of 2 μL using the syringe pump and attached cannula (n = 2 mice). Using a Nikon A1R multiphoton microscope with the same hardware as above, mice were imaged throughout the growth of the tumor using a combined tile and z-stack to capture the whole volume beneath the window. Analysis was performed using Imaris to create surfaces of the GFP (tumor cells) and mCherry signal at each time point and quantify the resulting tumor cell and mCherry volume.

### Focused ultrasound blood-brain barrier disruption with imaging for interstitial transport metrics

The focused ultrasound study was conducted at Virginia Polytechnic Institute and State University under approved protocol #21-072. 8-12 weeks old transgenic B6.Cg-S1pr3tm1.1Hrose/J mice (S1P_3_^mCherry^, The Jackson Laboratory #028624) were anesthetized with isoflurane and connected to a stereotactic frame. To serve as controls for a separate study, a burr hole was drilled at coordinates -2 mm, +1.2 mm, -1.5 mm (AP, ML, DV) from bregma. Mice were injected with 5 µL of sterile saline (0.9% NaCl) via a Hamilton syringe and syringe pump (World Precision Instruments) at 1 µL/min for 5 min. The Hamilton syringe was removed three minutes after the completion of injection to prevent reflux, and the burr hole was sealed with bone wax.

MRI parameters are described above. Following the pre-contrast baseline T1-weighted image, a bolus of 3.75ul/g mouse of microbubbles (OptisonTM, GE Healthcare #NDC 0407-2707-18, stock concentration 5x108 and 8x108 microbubbles/mL) was injected via a catheter (26G (BD), tubing length 50 cm, (PE50, BD427411)) into the tail vein. A magnetic-resonance imaging guided focused ultrasound system (RK-300; FUS Instruments Inc., Toronto, ON, Canada) with 2-axis (x-y) motorized spatial positioning was used to disrupt the BBB. Two targets were sonicated with coordinates A4.0, R2.0, and S4.2 for target #1 and A4.0, L1.1, and S4.2 for target #2. Ultrasound bursts (10 ms burst length; 1 s repetition period; 120 bursts per target location) were produced by a waveform generator (SIGLENT SDG1.32X) and amplified to drive a 1.15 MHz spherically-focused transducer (25 mm diameter, 20 mm radius of curvature, with central hole for 6 mm diameter passive detector receiver; FUS Instruments Inc.). This system has an integrated BBB feedback mode which ramps the pressure amplitude (in this case, from 0.15 to 0.715 MPa in 0.25 MPa steps) during the initial bursts and selects the optimal pressure amplitude based on the waveform received by the passive cavitation detector. This procedure resulted in amplitude of 0.34 MPa for target #1 and 0.36 MPa for target #2. The transducer was coupled to the mouse skull using ultrasound coupling gel and the axial offset (z) was adjusted manually to target the center of the brain. Immediately after sonication, contrast was delivered and T1-weighted images were collected over time.

### Tissue culture insert model

Hydrogels were seeded in 8 µm-pore diameter tissue culture inserts (Corning 3374). If needed, cells were fluorescently labeled with CellTracker dyes (Life Technologies) and Vybrant dyes (Life Technologies) according to the manufacturer suggested protocol. Glioblastoma cells including G34, G528, G2, and G62, (5.0 x 10^5^), astrocytes (6.25 x 10^4^), and microglia (6.25 x 10^4^) were seeded in 75 µL hydrogels, a ratio within the range of invasive GBM tissue, as published^5^. Briefly, cells were seeded in gel containing 2 mg/mL thiol-modified hyaluronan (Advanced Biomatrix GS220) and 1.2 mg/mL rat tail collagen I (Corning 354236) in serum-free medium (basal Astrocyte Medium, Sciencell 1801-b; 1% v/v B-27, Gibco 12587-010; 0.5% v/v N-2, Gibco 17502-048). Hydrogels were incubated at 37°C in a humidified incubator (5% CO_2_ and 21% O_2_) for 3 hours. Flow was generated via a 125 uL fluid pressure head (serum-free media) on crosslinked hydrogels, with 35 uL between the well-plate and insert membrane. Volumes were inverted for net static conditions. Drug treatments (S1PR3 inhibitor TY52156^30^; S1PR inhibitor CAY10444) were distributed at 10 μm in hydrogels prior to crosslinking, then applied to the external flow media. Controls were equal volumes of vehicle.

To quantify invasion, cell-laden gels were pipetted out of the tissue culture inserts and the membranes were fixed with 4% paraformaldehyde for 20 minutes at room temperature. The top of the membranes were carefully cleaned using small cotton tipped applicators. Cells that migrated through pores and attached to the opposite surface of the membrane were identified by CellTracker labels and DAPI staining. Three representative 10X fields were imaged per insert and averaged across technical replicates (individual gels) and reported as average cells invaded/total cells seeded x 100 (%).

### S1PR3 expression analysis under shear

Channels of an Ibidi µ-Slide VI 0.4 were coated with 1 mg/mL collagen in PBS for 30 minutes at 37°C, followed by washing in PBS. Astrocytes and microglia were independently labeled with CellTracker Green or Vybrant DiD intracellular dyes according to manufacturer instructions. Cell solutions of 300,000 cells/mL in complete Astrocyte Medium (Sciencell 1801) were added to the coated channels to plate 9,000 cells total. Channels were loaded with astrocytes only, microglia only, or both astrocytes and microglia (4,500 each). The device was incubated for 1 hour to allow for initial cell attachment, then 90 uL of complete medium was added to prevent dehydration during the overnight incubation. The following day, complete medium was exchanged for serum-free medium (basal Astrocyte Medium, Sciencell 1801-b; 1% v/v B-27, Gibco 12587-010; 0.5% v/v N-2, Gibco 17502-048) approximately 3 hours prior to the start of experiments.

Device inlets were attached to syringe pumps (Chemyx Fusion 200) using an Ibidi port adapter, and outlet ports were connected to open tubing for draining and collecting on the shelf below. For varying shear rate, the inlet flow rate was adjusted to achieve shear stresses of 0.01, 0.1, 0.5, or 1 dynes/cm^2^. For varied duration, the shear stress is held constant at a low pathological rate (0.1 dynes/cm^2^). Control channels were plugged with adapters to prevent dehydration but not connected to pumps (0 dynes/cm^2^). After the experiment, cells were immediately fixed in 4% paraformaldehyde for 20 mins at room temperature and washed with 1X PBS. Cells were fluorescently labeled for S1PR3 expression using the Alexa Fluor™ 555 Tyramide SuperBoost™ Kit (Invitrogen B40923). Three images from each well were taken at 20X magnification, and the number of cells are averaged per frame for each condition per trial (n = 4). We counted the total number of S1PR3^+^ cells and the number of S1PR3^+^ astrocytes or microglia. Data were reported as percentages of total cells/frame.

### In vivo invasion quantification

#### mCherry expression vs. invasion study in S1P ^mCherry^ mice

DAPI, GFP, and mCherry channels were split and exported individually using ImageJ. The GFP channel of GFP + GL261 cells was thresholded based on intensity to remove background fluorescence, and images were colocalized with DAPI staining to restrict analysis to GFP signals associated with nuclei. Watershed segmentation in ImageJ was applied to separate touching or overlapping nuclei, and the resulting objects were defined as individual GL261 cells. Control point image registration of GFP and mCherry IHC images was then performed in MATLAB using a local weighted mean geometric transformation to account for orientation and shape differences between MRI slices and histological tissue sections. For each registration, at least 12 distinct and reliable anatomical landmarks were selected on consistent gross anatomical features such as brain shape and ventricles to minimize modality-dependent bias. Finally, for each pixel on the MRI, the number of invading cells and the mCherry staining intensity were extracted, and the z-score–normalized mCherry brightness was calculated in pixels with and without invading cells.

#### TY52156 inhibitor study

Isograft tissue sections were stained and imaged for GFP as described above. Five standardized locations around the tumor border were imaged for each of three tissue sections. False-colored images were compiled in ImageJ, and the tumor border was visually demarcated based on DAPI staining. The number of tumor cells invaded past the tumor border were counted per frame, normalized to frame coverage area (cells/mm^2^), and averaged for each animal.

### Convection enhanced delivery

One week after intracranial tumor implantation, tumor-bearing mice were re-positioned on the stereotactic frame and the original burr hole was re-exposed. A blunt-end needle was attached to a Hamilton syringe and syringe micropump (UMP3, World Precision Instruments) and inserted 2.2 mm below the surface of the skull. Sterile saline was infused at 1 uL/min for 10 minutes. The needle was left in place for 3 minutes to prevent reflux, then the needle was slowly removed. The burr hole was refilled with dental wax, and the animal was allowed to recover for two days before tissue collection.

### Rotational shear model

50,000 cells per well of astrocytes and/or microglia were seeded on collagen I-coated 6-well plates in complete Astrocyte Medium (Sciencell 1801) and attached overnight at 37°C 5% CO_2_ in a humidified incubator. The following day, serum-containing medium was removed and replaced with serum-free medium (basal Astrocyte Medium, Sciencell 1801-b; 1% v/v B-27, Gibco 12587-010; and 0.5% v/v N-2, Gibco 17502-048). Well plates for shear conditions were placed on an orbital shaker for 18 hours at 100 RPM (VWR Orbital Mini Shaker 444-0268) in the incubator. With a 9.5 mm radius of rotation, this is expected to subject cells to ∼1 dynes/cm^2^ of shear^63^. Rest controls were placed on the adjacent shelf.

### Conditioned media generation

#### 2D glial cell conditioning

50,000 astrocytes, microglia, or both were plated in collagen-coated 6-well plates in Astrocyte Complete Medium (2% serum) and allowed to adhere overnight. Approximately 3 hours before the start of shear, the cells were serum-starved in serum-free medium (basal Astrocyte Medium, Sciencell 1801-b; 1% v/v B-27, Gibco 12587-010; 0.5% v/v N-2, Gibco 17502-048). For blocking studies, CAY10444 was added at 10 μM in DMSO during the serum-starve phase. Inclusion of TY52156 in a similar manner (10 μM) resulted in glial cell death, hence the use of CAY10444 instead. The sheared plate was transferred to an orbital shaker (VWR Orbital Mini Shaker 444-0268) at 100 RPM, producing an estimated shear stress of 1 dynes/cm^2^. Rest controls were placed on the adjacent shelf. Conditioned media was collected 18 hours after shear and either used immediately or stored at -20°C until use. Sequential shear conditioned medium was collected from the first sample (e.g., sheared astrocytes) and immediately transferred to the second sample (e.g., static microglia), to be collected a final time the following day (i.e., two days of conditioning).

### Chemotaxis experiments

To assess glioma cell chemotaxis, 50 uL of growth factor reduced basement membrane extract (2 mg/mL; Trevigen) was plated into each well of a 96-well tissue culture insert and allowed to gel at 37°C for 30 minutes. Once gelled, 125 uL of control or conditioned medium was added to the bottom of the well, and 50,000 GSCs are plated on top of the hydrogel in serum-free medium (basal Astrocyte Medium, Sciencell 1801-b; 1% v/v B-27, Gibco 12587-010; and 0.5% v/v N-2, Gibco 17502-048). The well plate was placed back in the incubator for 18 hours.

### Database analyses on bulk GBM

Bulk tumor (TCGA-GBM) versus postmortem brain tissue (GTEx) S1PR3 RNAseq comparisons were made using the GEIPA2 database within the online analysis program using a | Log_2_FC | cutoff of 1 and p-value cutoff of 0.01^31,64^. The GEIPA database matches TCGA tumor samples to non-diseased GTEx samples following review by medical experts for sample grouping^31^. Raw expression data is recomputed based on a standardized pipeline and is provided by the UCSC Xena Project (http://xena.ucsc.edu/)^31^. The GTEx dataset provides comparison tissues from postmortem, non-diseased donors based on a series of exclusion criteria and pathology review^65,66^.

cBioPortal is a public database site developed by the Memorial Sloan Kettering Cancer Center, compiling cancer datasets across pathologies^33–35^. We queried the Clinical Proteomic Tumor Analysis Consortium dataset^32^ for S1PR3 proteomic data in bulk GBM generated by LC-MS/MS. A major advantage of this dataset is that it includes information pertaining to patient demographics and outcomes (de-identified). High- and low- S1PR3 expressing patients were grouped via the online portal analysis interface based on protein abundance ratio z-scores ± 1. S1PR3 protein abundance ratio, S1PR3 protein abundance ratio Z-scores, S1PR3 high-low assignment, days from diagnosis to death, and days from diagnosis to recurrence values were exported for plotting and analysis in GraphPad Prism 10.2.0.

### Immunohistochemistry in patient tissue

#### GBM invasive front tissue microarray

The GBM invasive edge TMA was constructed by the Tumor Tissue and Pathology Shared Resource of the Wake Forest Baptist Comprehensive Cancer Center with the assistance of the Wake Forest Brain Tumor Center of Excellence. Hematoxylin and eosin-stained sections from all FFPE tissues were reviewed by a pathologist to demarcate the tumor/brain interface. One 2 mm tissue punch was obtained from the tumor/brain interface in each specimen block. FFPE tissue cores from 42 GBM patients (average age 67.7 ± 10.9 years, 25 male, 17 female) were included in the TMA. Sections were cut at 5 uM and mounted on positively charged slides. Tissue was stained using the custom α-S1PR3 N-terminal antibody (Aves Labs) described above. Chromogenic IHC was performed using Vectastain Elite ABC reagents (Vector Labs) using the peroxidase substrate DAB and counter-stained with hematoxylin. Tissue cores containing any number of S1PR3^+^ cells were considered positive. S1PR3^-^ cores had no S1PR3^+^ cells. Cell counts for the phenotypic markers for neurons and glial cells, the neuroinflammation markers and immune cells were compared between S1PR3^+^ and S1PR3^-^ cores.

#### GBM and brain-metastatic tissue

Patient samples were accessed through the University of Virginia Biorepository and Tissue Research Facility. GBM patient samples were selected by a neuropathologist (J. Mandell) based on a definitive diagnosis of glioblastoma (astrocytoma, World Health Organization grade IV) who had completed tumor resections at the University of Virginia between 2010 and 2013. Samples were de-identified and processed to select tumor sections that included a portion of adjacent non-bulk tumor tissue (here referred to as the parenchyma interface) as identified by a neuropathologist (F. Bafakih). Formalin fixed paraffin embedded 8 µm sections were deparaffinized with xylene and rehydrated in graded ethanol, treated with high pH antigen unmasking solution (Vector Labs), and stained with anti-S1PR3 (R&D Systems) followed by DAB substrate (Vector) according to manufacturer suggested protocols and counterstained with hematoxylin (Thermo Scientific). Hematoxylin and eosin staining was performed by the University of Virginia Biorepository and Tissue Research Facility following standard protocols. Areas at the tumor-parenchyma invasive front of tumor resections were imaged using wide-field microscopy with EVOS FL Auto (Life technologies) and Aperio Scanscope (Leica Biosystems) and quantified using ImageJ (National Institutes of Health).

### qRT-PCR

Tissues were flash-frozen at the time of collection and stored at -80°C until RNA was extracted. For RNA extraction, frozen tissues were dropped into a Dounce homogenizer prefilled with 1 mL TRIzol (ThermoFisher 15596026) and homogenized 10-15 strokes. Chloroform (200 µL, Millipore Sigma C2432) was added, samples were mixed and incubated for at least 5 min and centrifuged at 4°C to generate the RNA-containing aqueous layer (∼600 µL). One mL of 100% ethanol was added to the aqueous layer and the RNA was purified using the Quick-RNA Miniprep kit (Zymo Research R1054) following the manufacturer’s protocol including the DNAse treatment. RNA was quantified with a Nanodrop Lite (ThermoFisher ND-LITE-PR). Only samples with a 260/280 ratio of 1.9-2.1 were used in the RT-PCR analysis. Total RNA (100 ng/sample) was reverse transcribed with the QuantiTect Reverse Transcription kit (Qiagen 205311) following the manufacturer’s protocol. A standard curve was generated by making a pool of all the samples and performing RT reactions ranging from 30 ng to 500 ng of RNA allowing quantification of the samples by comparison to this standard curve. A RT-minus reaction was also included as a control. PCR was performed with TaqMan assays (S1PR3 Hs01019574_m1, housekeeping ATP5F1 Hs01076982_g1, ThermoFisher) and TaqMan Universal Master Mix II (ThermoFisher 4440047) following the manufacturer’s directions. PCR reactions were run on a LightCycler 96 (Roche 5815916001). Calculated S1PR3 expression was normalized to ATP5F1 expression. Descriptive statistics, unpaired t-test and correlations were performed using GraphPad Prism version 10.0.0 for Windows, GraphPad Software, Boston, Massachusetts USA, www.graphpad.com.

### Human magnetic resonance imaging for interstitial transport metrics and image-localized biopsies

Human studies were performed at Carilion Clinic in Roanoke, VA, on Protocol 22-1590 approved by the Carilion Clinic Institutional Review Board. For human imaging, DCE-MRI was performed on patients with suspected glioblastoma. Patients were imaged with a 3.0 T (SIEMENS MAGNETOM 3.0T XQ Numaris/X VA60A-0CT2) equipped with maximum gradient amplitudes of 80 mT/m and slew rates of 200 T/m/s. T2-weighted images were acquired to visualize anatomical features and tumor location. T2 parameters were as follows: repetition time (TR) = 4000 ms, echo time (TE) = 84 ms, field of view (FOV) = 263 x 350 x 350mm with a 264 x 320 matrix, slice thickness = 3 mm, number of slices = 32. A pre-contrast T1-weighted image was collected for baseline signal normalization. Then, gadolinium (Magnevist, Bayer HealthCare Pharmaceuticals) was delivered and multiple post-contrast T1-weighted images were acquired over time. T1 parameters were: n post-contrast T1 = 52, TR = 9.3 ms, TE = 4.29 ms.

### nanoString analysis on patient GBM

Patient GBM biopsies were submitted to Nanostring for processing with the nCounter® Neuroinflammation Panel, designed to calculate the abundance of CNS-resident and peripheral immune cells^67^. For quantifying cell type abundance, data was imported into ROSALIND® (https://rosalind.bio/). Data was analyzed with a HyperScale architecture developed by ROSALIND, Inc. (San Diego, CA). Read Distribution percentages, violin plots, identity heatmaps, and sample MDS plots were generated as part of the QC step. Normalization, fold changes and p-values were calculated using criteria provided by Nanostring. ROSALIND® follows the nCounter® Advanced Analysis protocol of dividing counts within a lane by the geometric mean of the normalizer probes from the same lane. Housekeeping probes to be used for normalization are selected based on the geNorm algorithm as implemented in the NormqPCR R library1^68^. Abundance of various cell populations is calculated on ROSALIND using the Nanostring Cell Type Profiling Module. ROSALIND performs a filtering of Cell Type Profiling results to include results that have scores with a p-Value greater than or equal to 0.05. Fold changes and pValues are calculated using the fast method as described in the nCounter® Advanced Analysis 2.0 User Manual. P-value adjustment is performed using the Benjamini-Hochberg method of estimating false discovery rates (FDR). Clustering of genes for the final heatmap of differentially expressed genes was done using the PAM (Partitioning Around Medoids) method using the fpc R library2 that takes into consideration the direction and type of all signals on a pathway, the position, role and type of every gene, etc.^69^. Hypergeometric distribution was used to analyze the enrichment of pathways, gene ontology, domain structure, and other ontologies^70,71^. The topGO R library3^72^, was used to determine local similarities and dependencies between GO terms in order to perform Elim pruning correction. Several database sources were referenced for enrichment analysis, including Interpro4^28^, NCBI5^71^, MSigDB6,7^73^, REACTOME8^74^, WikiPathways9^75^. Enrichment was calculated relative to a set of background genes relevant for the experiment. Gene expression heatmaps were sorted using the ratio of the peritumoral biopsy with the most tumor originating pathlines (B3) over the geometric mean of the other peritumoral biopsies (B1 and B2).

### Statistics

Descriptive statistics, unpaired t-tests, two-tailed t-tests, correlations, and Kaplan-Meier analyses were performed with GraphPad Prism 10.0.0-10.2.0 for Windows (GraphPad Software, Boston, Massachusetts USA, www.graphpad.com) with *p* < 0.05 considered significant. For experiments, statistics were performed across at least 3 biological replicates (i.e., a single animal or an independent experiment/trial) across multiple cell passages where relevant. Illustrations were created with BioRender (www.biorender.com).

## Supporting information

Supplemental Figures with Legends

## Acknowledgements

Michelle Olsen, Kevin Lynch, Jony Kipnis, Mark Okusa, Amandeep, Amy Rizzo, BTRF, Molecular biology core UVA, Jordan Darden, Amy Wahba, Jack Roy, Jason Raymond, Flow Cytometry Core, Biocomplexity Institute, ICTAS, Shayn Peirce-Cottler, Jessica Boni, Tristan Stoyanof, Savieay Esparza, BTRF and the Wake Forest pathology core

