## Supplemental Figures with Legends for "S1PR3 mediates glial stimulated tumor invasion in response to interstitial fluid flow"

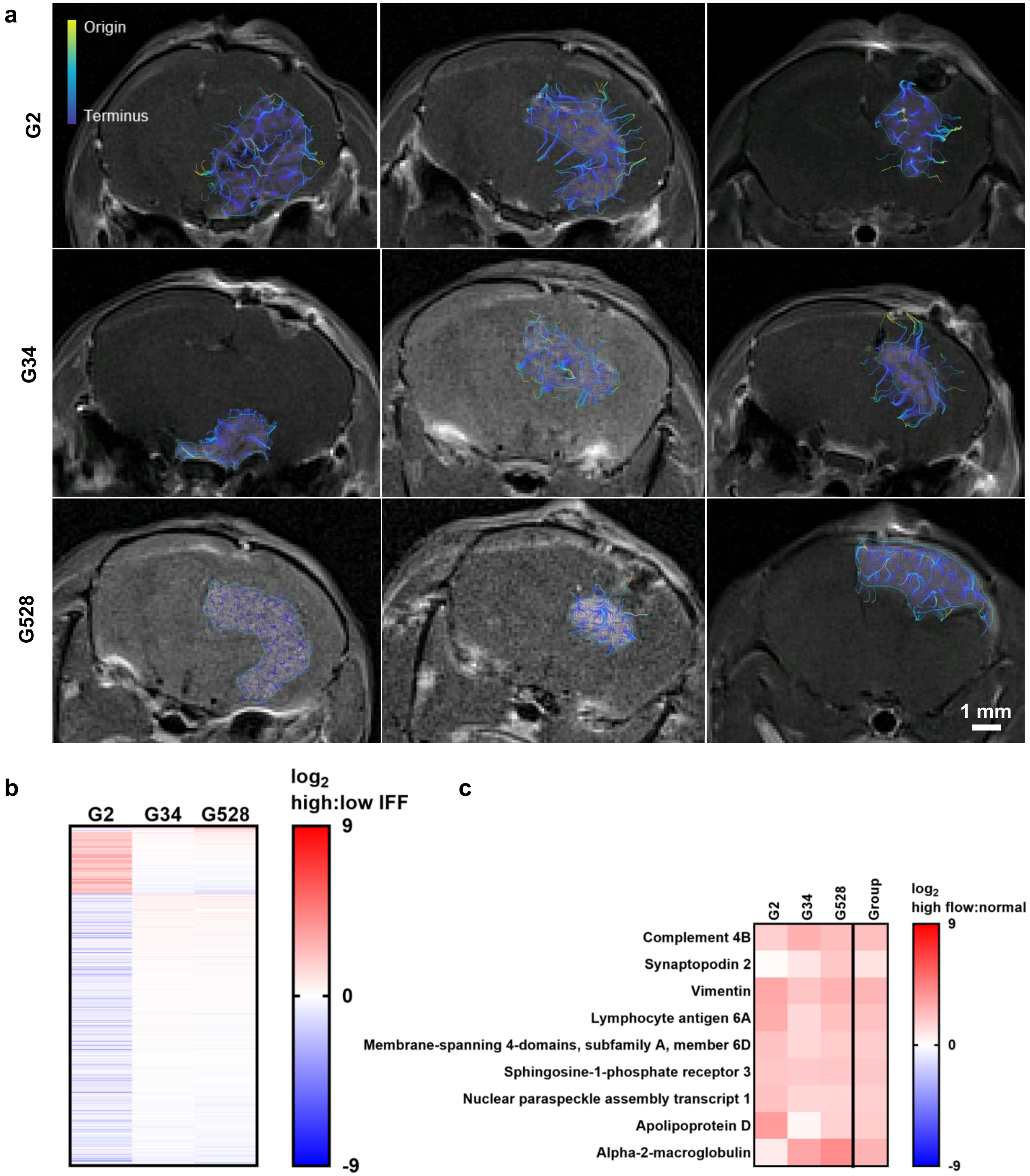


**Extended Figure 1. Full microarray expression data in high vs. low peritumoral flow regions and comparison with bulk tumor and normal tissue. a.** PDXs were imaged with DCE-MRI at 11 days post-implantation to calculate interstitial transport metrics. Tumor originating pathlines are calculated from the resulting flow magnitude vector field and are shown. **b-c**. Biopsies from Evan's blue-indicated high and low flow regions adjacent to the tumor were processed for microarray against neuroinflammatory genes (ArrayST 2.0) and are sorted by mean normalized gene expression values across the PDXs. The full gene panel normalized to low flow peritumoral regions is shown in **b**. For comparison, the same top 9 hits are shown normalized to non-transformed tissue in (**c**).

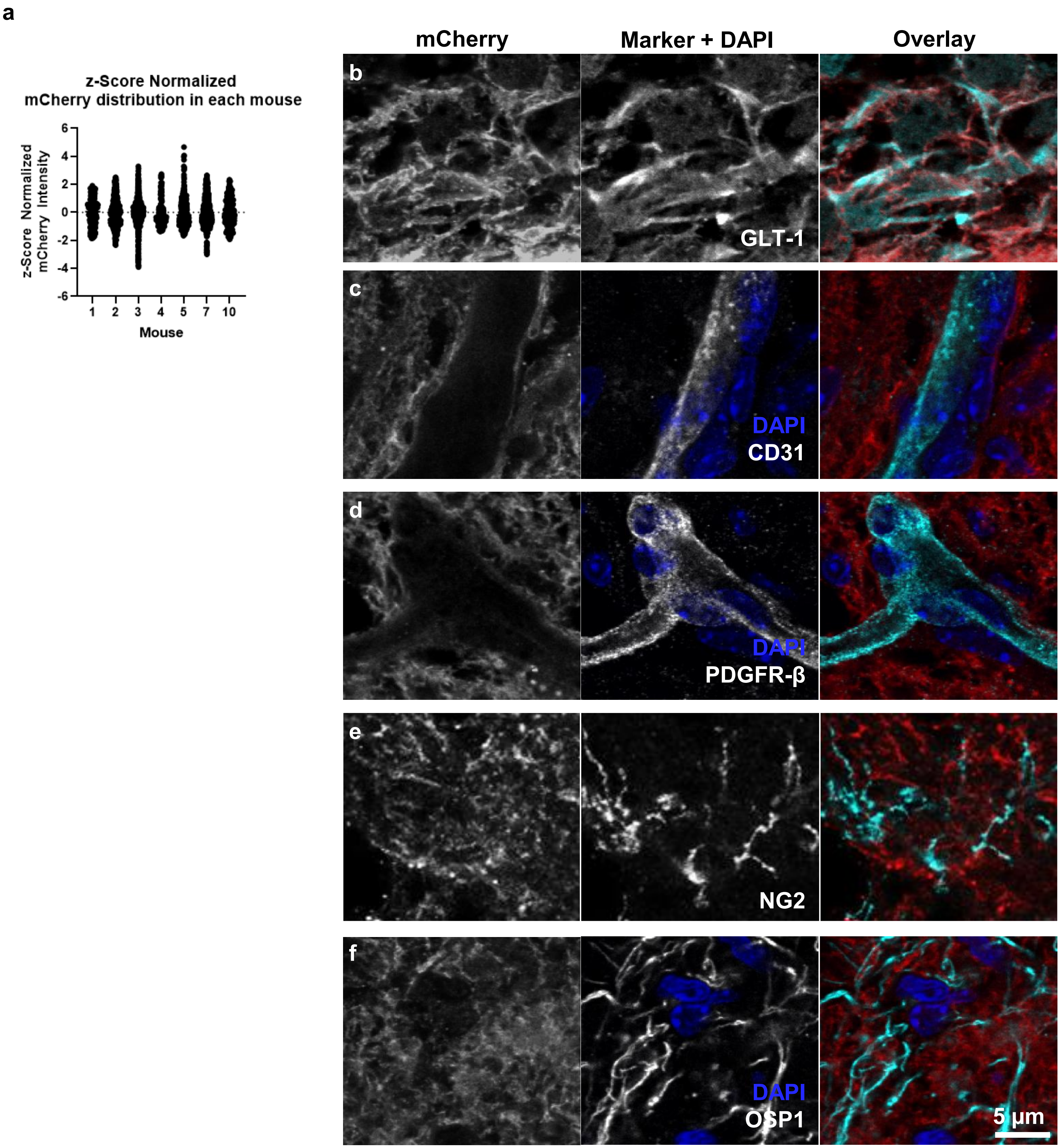


**Extended Figure 2. mCherry expression co-localizes with markers of astrocytes (GLT-1) in outward flow regions but not with other cell markers. a.** S1PR3$^{mCherry}$ mice were implanted with GFP-GL261 and underwent MRI on post-implant days 15 or 24. Tissue was collected one day post DCE-MRI and tissue sections were composite (n = 3 sections) and registered to MRI. MRI-pixel distribution of reporter fluorescence intensity is plotted for each mouse. **b.** Immunofluorescence across cell-type markers in S1PR3$^{mCherry}$, GFP-GL261 bearing mice.

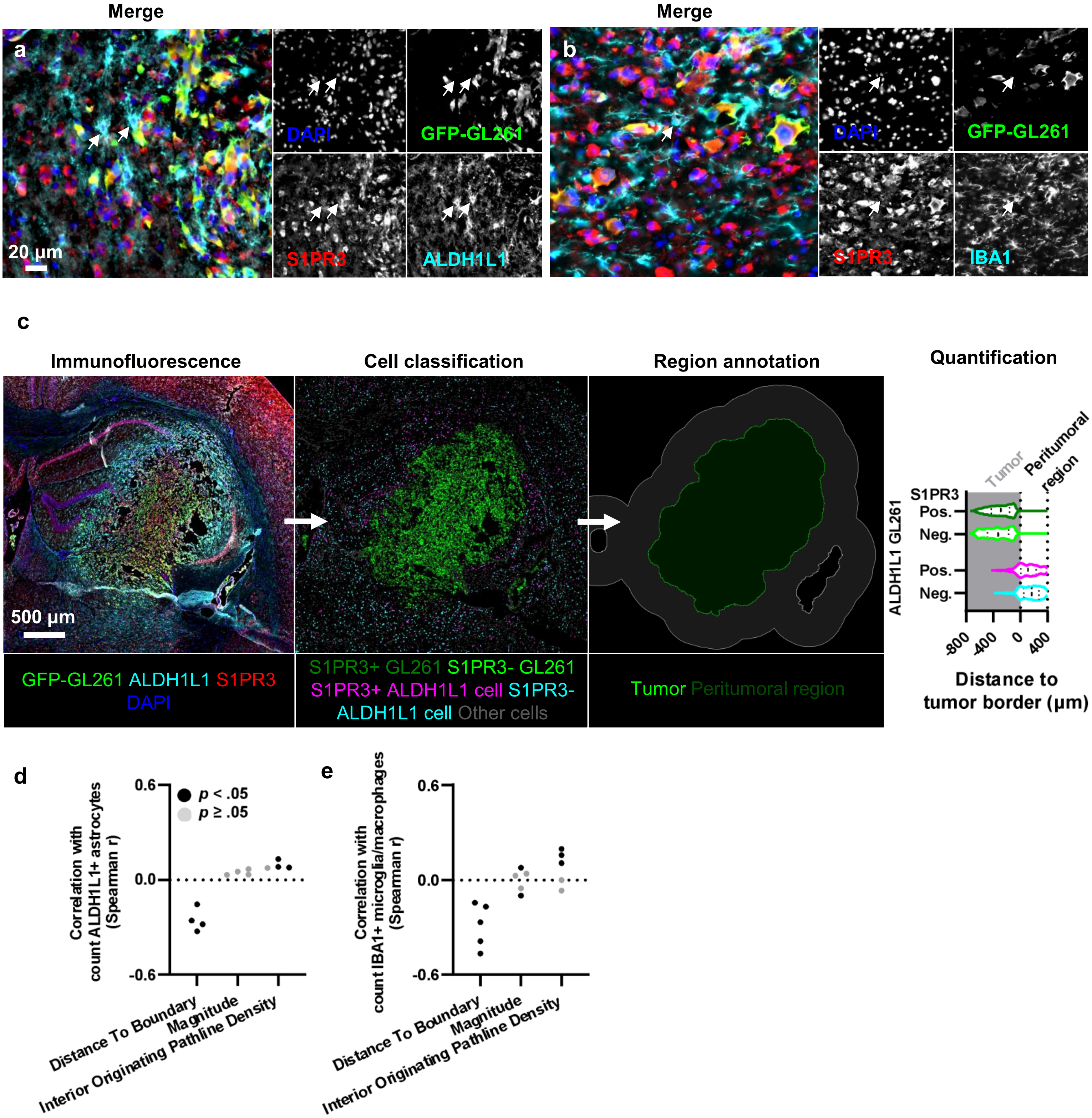


**Extended Figure 3. S1PR3 expression in glia exhibits a specific spatial distribution in the peritumoral region. a-b.** Immunofluorescence for S1PR3 and ALDH1L1 (astrocytes, **a**) or IBA1 (microglia/macrophages, **b**) in GFP-GL261 tumors. **c**. Cell classifications generated from a random-forest model in QuPath. A threshold of the density of GFP-GL261 (S1PR3+ and -) classifications defined the tumor border for distance to tumor border calculations (cells within 400 µm of the border). **d.** Correlation between counts of total classified cell types (astrocytes, **a**; microglia **e**) and flow metrics (n = 4-5 mice per stain).

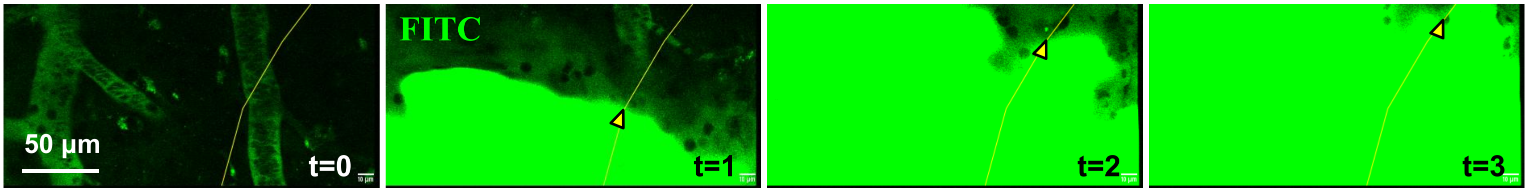


**Extended Figure 4. Flow rate analysis of dextran through cannula**. A tumor-naïve S1P3$^{mCherry}$ mouse was fitted with an optical window and the brain parenchyma was infused with 3 uL/min FITC-dextran via an intracranial catheter. The the leading edge of FITC-dextran was tracked (yellow arrow and line). This pump infusion rate resulted in 12 µm/s interstitial fluid velocity.

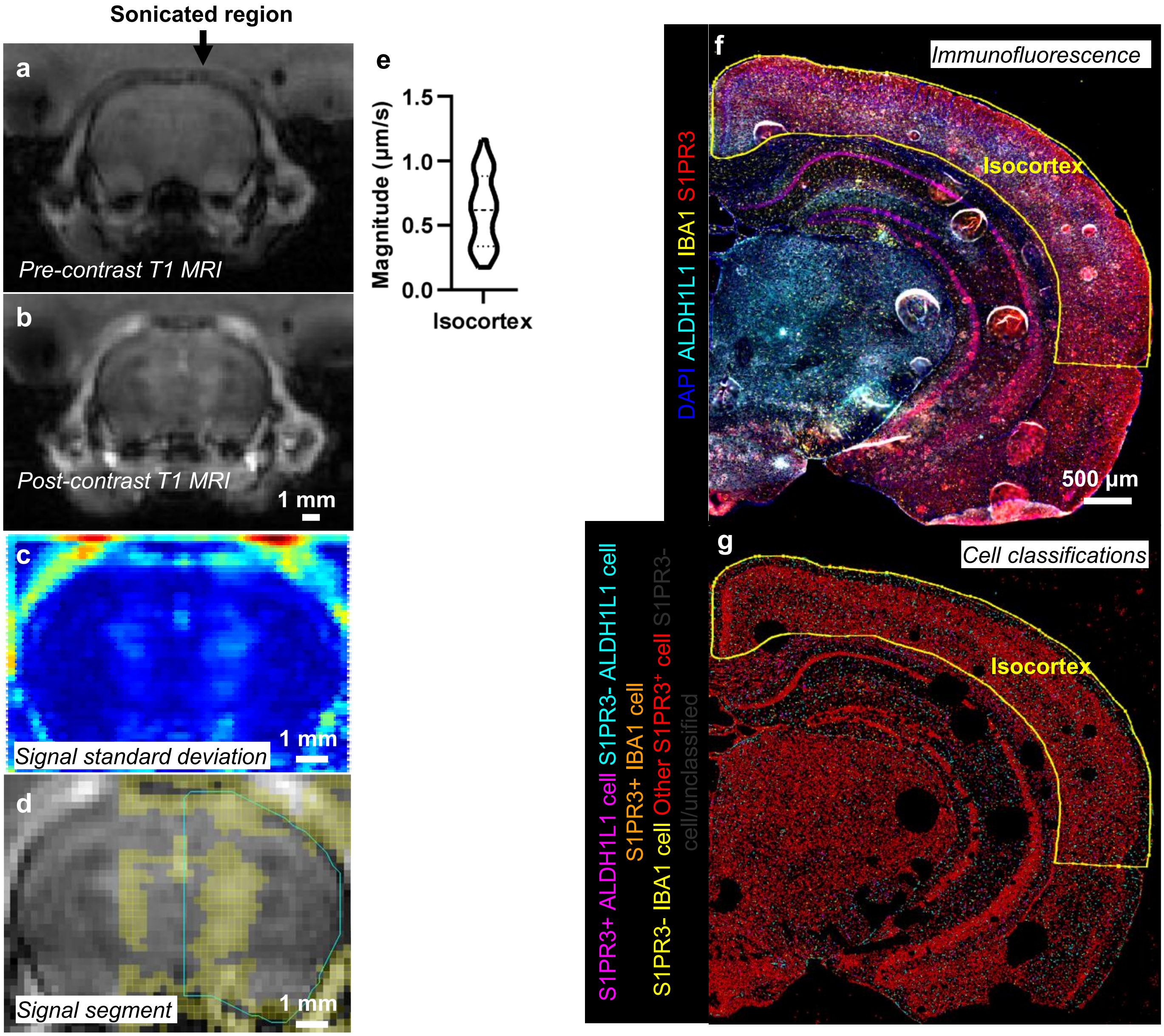


**Extended Figure 5. Transport metric quantification and cell classification schemes for sonicated tumor-naïve mice. a-b.** DCE-MRI was performed to quantify interstitial transport metrics immediately post- focused-ultrasound mediated blood-brain barrier disruption, showing pre-contrast T1-weighted MRI (**a**) and first post-contrast image (**b**). **c-d.** Contrast signal standard deviation (**c;** low blue; middle yellow-green; high red) was used to segment (**d**) the enhanced region for calculating interstitial transport metrics. **e.** Distribution of pixel-by-pixel interstitial flow velocity magnitude in the isocortex. **f.** Two days post-sonication, tissue was collected for immunofluorescence staining against ALDH1L1 (astrocytes), IBA1 (microglia/macrophages), and S1PR3. The isocortex was segmented by hand based on anatomical references. **g.** Cell classifications defining astrocytes, microglia, and bindary S1PR3 expression were defined in QuPath using a Random Forest model.

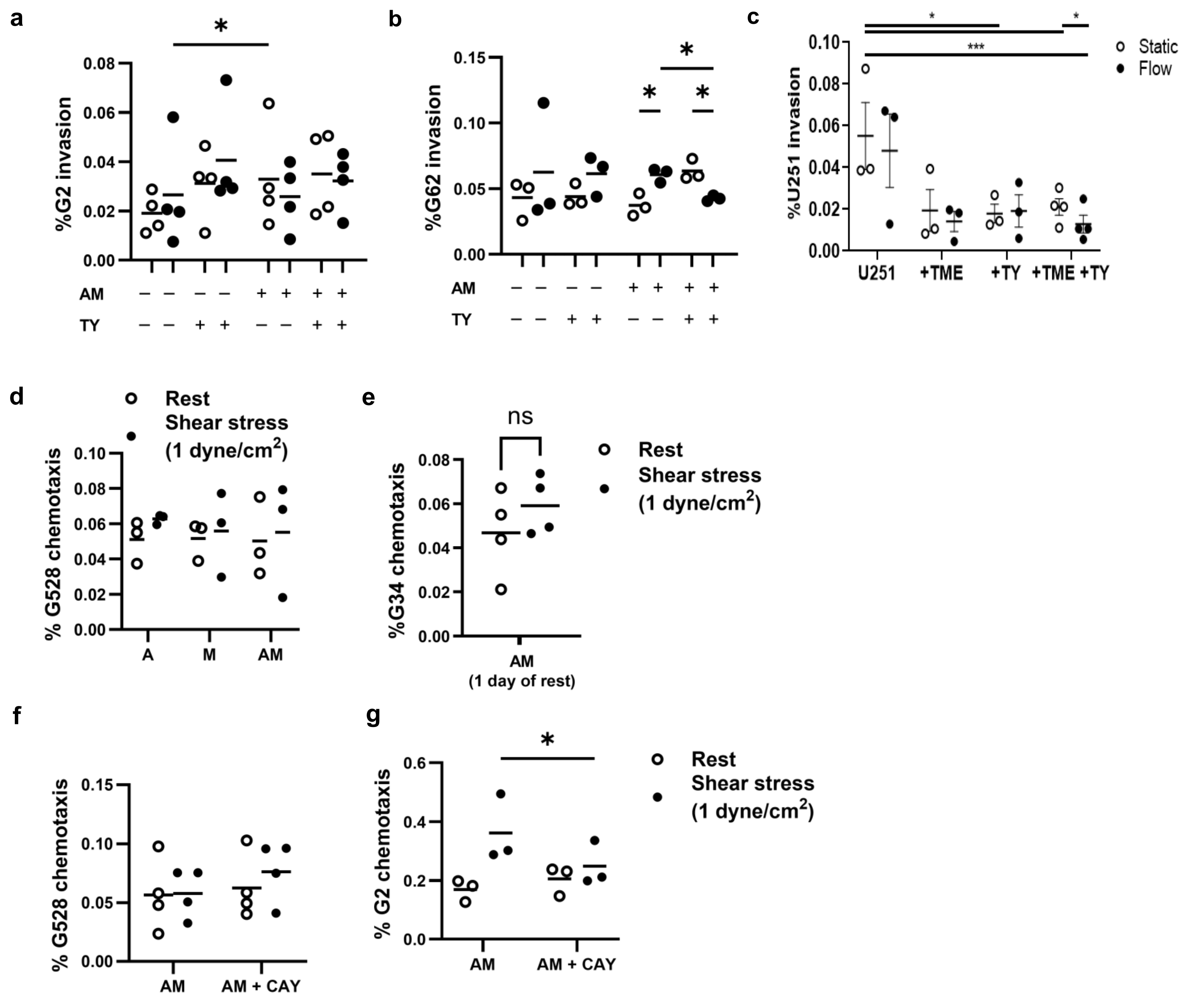


**Extended Figure 6. Blocking S1PR3 reduces invasion and chemotaxis for multiple GSC lines. a-c.** TME model containing astrocytes, microglia, and tumor cells in a 1.2 mg/mL collagen, 2.0 mg/mL hyaluronan hydrogel on a porous membrane to permit invasion over 18-20 h. Gels are placed under a fluid pressure head to permit flow through the matrix. G2 (**a**), G62 (**b**), and U251 (**c**) invasion demonstrating effect of flow, glia, and 10 µM TY52156 (n = 3-4 independent experiments/GSC line; RM one-way ANOVA with Tukey's multiple comparisons in **a** and Fisher's LSD in **b**). **d-g**. Chemotaxis model where mono- or co-cultured glia are sheared 18-20 h at 1 dyne/cm$^2$, then conditioned media is added to a well plate below a tissue culture insert containing GSCs suspended in the hyaluronan-collagen gel. GSCs are allowed to invade through the porous membrane toward conditioned media for 18 h. **d.** Chemotaxis model with G528 exposed to shear-conditioned media from isolated or co-cultured glia (n = 3 independent experiments; RM two-way ANOVA with Tukey's multiple comparisons). **e.** G34 chemotaxis model using conditioned media from sheared co-cultured glia collected one day post-shear (n = 4 independent experiments; ratio paired t-test). **f-g.** G528 (**f**) and G2 (**g**) chemotaxis models using conditioned media from sheared co-cultured glia in the presence of the S1PR inhibitor CAY10444 (n = 4 independent experiments; RM two-way ANOVA with Fisher's LSD). A, astrocytes; M microglia; *$p < .05$; ***$p < .001$.

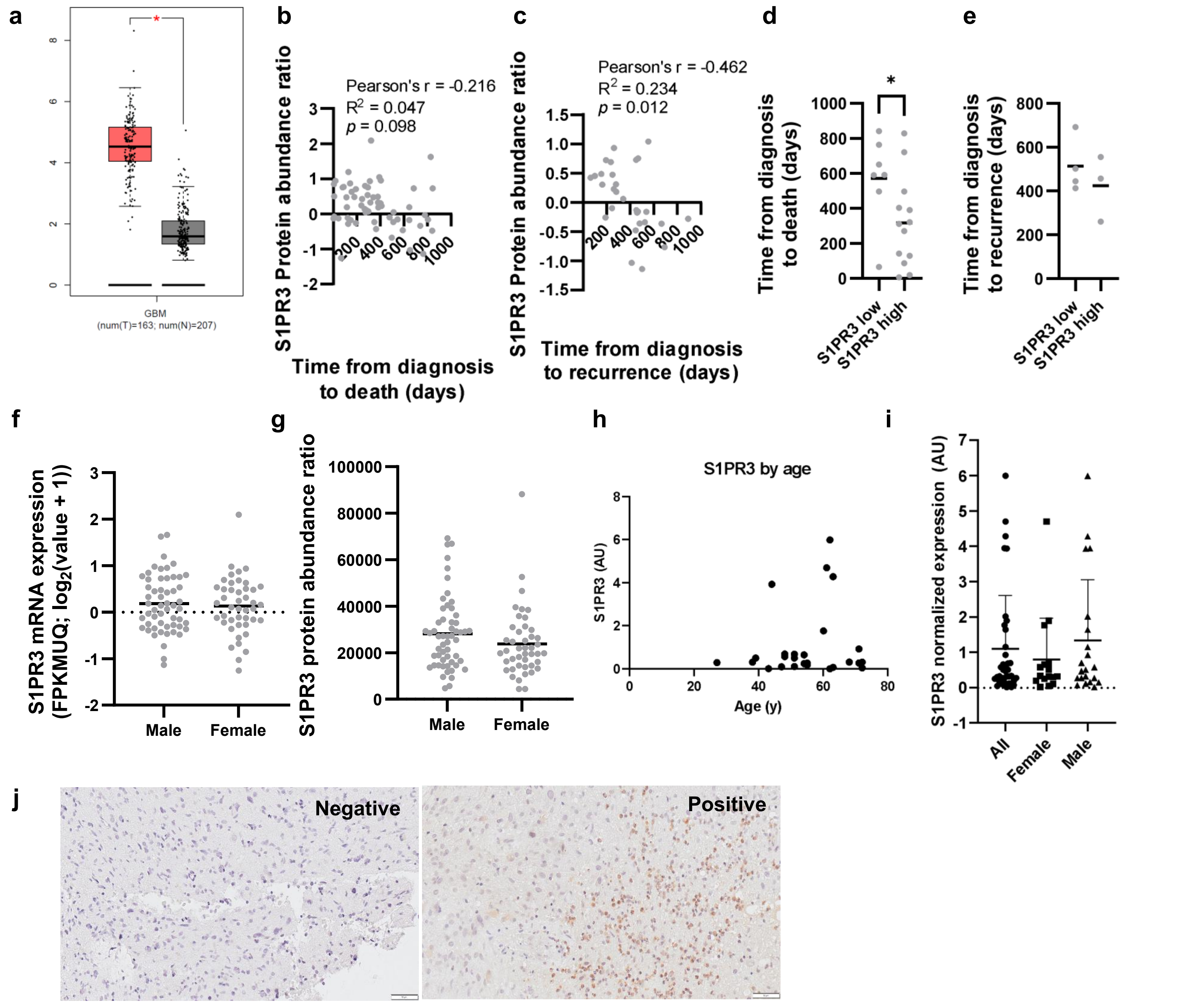


k

| | |
|---|---|
| n negative S1PR3 | 13 |
| n positive S1PR3 | 21 |
| **Total** | **34** |

**Extended Figure 7. Survival and demographic correlates with S1PR3 in bulk tumor and invasive front. a.** Bulk GBM tumor vs. postmortem brain RNAseq available from the GEIPA2 database (n = 163 tumor, 207 normal). **b-c**. Mass spectroscopy data collected by the Clinical Tumor Proteomic Analysis Consortium (CPTAC) and accessed via cBioportal (n = 29 patients) correlated with time from GBM diagnosis to death (**b**) and recurrence (**c**). **d-e.** CPTAC mass spectroscopy dataset grouped into high- and low-S1PR3 expressing patients by protein abundance z-scores ± 1 relative to the full dataset, showing time from GBM diagnosis to death (**d**) and recurrence (**e**) (t-test). **f-g**. RNAseq mRNA expression (**f**) and mass spectroscopy protein abundance (**g**) from CPTAC bulk GBM dataset (n = 96 patients) across sexes (t-test not sig.). **h-i**. RT-qPCR against from bulk GBM tumor samples compared to age (**h**) and sex (**i**) (n = 26 patients). **j-k.** A patient tissue microarray (TMA) was generated from GBM biopsies and cores representing invasive regions were pathologist-selected for TMA assembly. Chromogenic staining on TMA for S1PR3 (**j**) and count cores containing at least one S1PR3 positive cell versus none (n = 34 cores/patients). *p < .05.

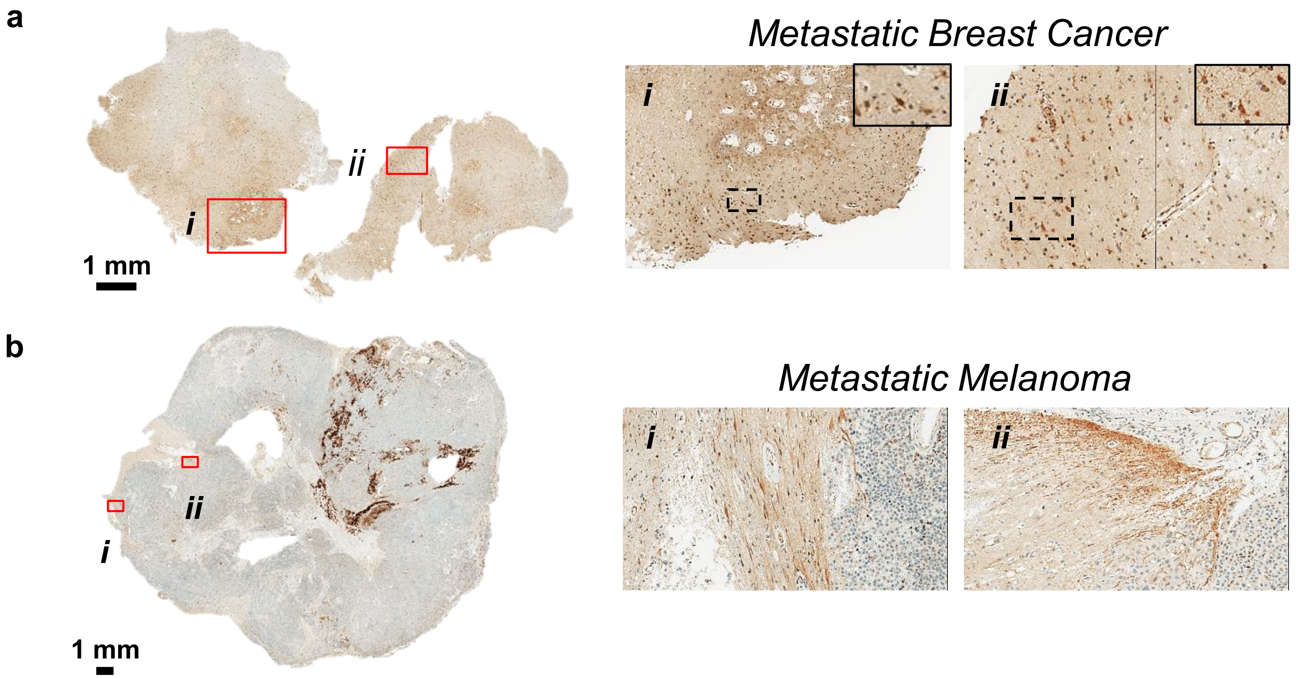


**Extended Figure 8. Positive S1RP3 expression in tissue cores from metastatic brain cancer.** Chromogenic staining for S1PR3 in patient tissues of breast-to-brain (**a**) and melanoma-to-brain (**b**) metastases.

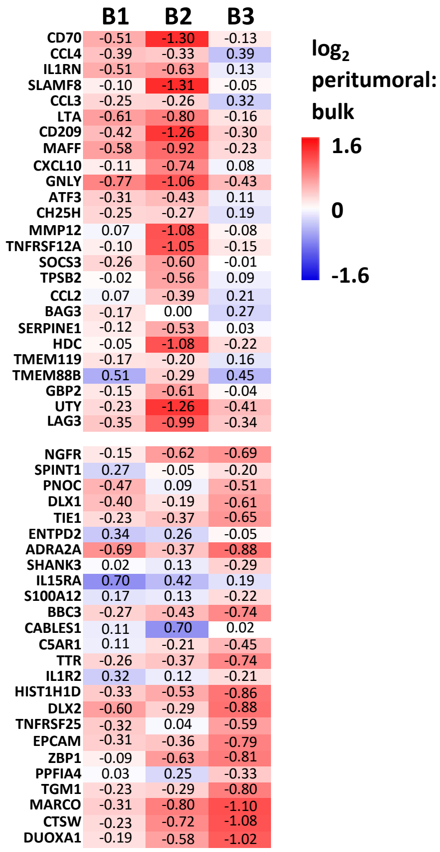


**Extended Fig. 9. Fold-change gene expression between peritumoral and bulk tumor from MRI-localized biopsies.** MRI-guided biopsies collected for S1PR3 RT-qPCR and neuroinflammation microarray designed to identify CNS-resident and peripheral immune populations and relevant genes. Top- and bottom-25 hits are shown, sorted by highest expression in B3 (highest n tumor originating flow pathlines) relative to the geometric mean of B1 and B2.
